# spVelo: RNA velocity inference for multi-batch spatial transcriptomics data

**DOI:** 10.1101/2025.03.06.641905

**Authors:** Wenxin Long, Tianyu Liu, Lingzhou Xue, Hongyu Zhao

## Abstract

RNA velocity has emerged as a powerful tool to interpret transcriptional dynamics and infer trajectory from snapshot datasets. However, current methods fail to utilize the spatial information inherent in spatial transcriptomics and lack scalability in multi-batch datasets. Here, we introduce spVelo (**sp**atial **Velo**city inference), a scalable framework for RNA velocity inference in multi-batch spatial transcriptomics data. Our model comparison studies show that spVelo compares favorably to existing methods regarding velocity consistency, transition accuracy, and direction correctness across expression levels, spatial graphs in each batch, and MNN graphs between batches. Furthermore, spVelo supports several downstream applications, including uncertainty quantification, complex trajectory pattern discovery, biologically significant state driver marker identification, gene regulatory network inference and temporal cell-cell communication inference. In conclusion, spVelo has the potential to provide deeper insights into complex tissue organization and underscore their biological mechanisms based on spatially-resolved patterns.

## 1 Introduction

Advances in sequencing technology have facilitated the reconstruction of cellular trajectories, revealing underlying dynamic processes [1–3]. Trajectory inference methods typically order cells along the pseudo-time axes based on similarities in their expression patterns [4–7]. However, traditional trajectory inference methods usually require prior knowledge of initial states or rely on certain assumptions, limiting the reliability and interpretability of these methods [5].

Recently, RNA velocity has become an alternative approach for trajectory inference. RNA velocity describes the rate of expression change for a single gene at a given time point, based on spliced and unspliced counts of messenger RNA (mRNA) [8]. The velocities of genes can then be used to estimate the future transcriptional states of cells, offering a powerful tool for understanding cellular differentiation, lineage tracing, and dynamical processes [9].

Current popular RNA velocity methods make different modeling assumptions. Velo-cyto [8] used a steady state and a linear relationship between unspliced and spliced RNA counts to model transcriptomics data. scVelo [10] relaxed this assumption by modeling cell-specific&gene-shared latent time and gene-specific kinetic rates. The stochastic mode of scVelo uses both first- and second-order moments and solves the steady-state ratio using generalized least squares. The dynamical mode of scVelo models kinetic rates, latent time, and transcriptional state as latent parameters, and fits linear differential equations with EM algorithms for each gene. Complex dynamics may violate the linear assumptions made by scVelo [11]. To solve this problem, UniTVelo [12] modeled spliced gene expression using radial basis function (RBF) instead of ODEs, allowing more flexible gene expression profile modeling. LatentVelo [13] utilized neural ordinary differential equations (neural ODEs [14]) on embedded latent space while performing batch effect correction. The annotated mode of LatentVelo further adds cell type information by modifying the prior. veloVI [15] is a Bayesian deep generative model using a gene-shared latent variable for summarizing the latent state of each cell.

While these methods have been successfully used to infer cellular dynamics [16, 17], they also suffer from several limitations [11, 18]. For example, current RNA velocity inference methods are confined to scRNA-seq data, which only captures the transcriptional profiles, losing the spatial context [19]. Spatial transcriptomics, a rapidly emerging technology, addresses this limitation by measuring the spatial information of gene expression. Spatial resolution determines the relative positions of cells and further reflects the communication and transitory relationships between adjacent cells. Utilizing spatial information can enable better inference of RNA velocity and trajectory, proven by the ablation test in Extended Data Fig. 1. Furthermore, current methods are confined to velocity inference in a single batch. This prevents the methods from utilizing the information from the entire dataset, thus failing to capture the global dynamics. Finally, current methods suffer from strict modeling assumptions about transcriptional kinetics. This assumption may not hold in complex biological systems where kinetic parameters can vary substantially among different genes, leading to poor RNA velocity inference in complicated dynamical features such as transcriptional boost [20], lineage-dependent kinetics and weak unspliced signals [11].

To address these limitations, we present spVelo (**sp**atial **Velo**city inference), a method for estimating RNA velocity in multi-batch spatial transcriptomics data. spVelo combines a Variational AutoEncoder (VAE) [21] for gene expression data with a Graph Attention Network (GAT) [22] for spatial location. By further adding a Maximum Mean Discrepancy (MMD) penalty [23] between latent spaces of different batches, spVelo is able to perform RNA velocity inference in a multi-batch spatial dataset. We compare spVelo with alternative methods using spatial data simulated from mouse pancreas data [24] and real oral squamous cell carcinoma (OSCC) data [25]. spVelo outperforms the previous RNA velocity inference methods for inferring RNA velocity and trajectory. Then, we demonstrate spVelo’s ability to perform batch effect correction on RNA velocity [13]. By leveraging the distributions of latent space, spVelo is able to quantify the uncertainty of the inferred latent state. We further show that spVelo can discover complex trajectory patterns, while other methods tend to predict a linear trajectory between cell types. By visualizing predicted phase portraits, spVelo is able to fit the genes’ dynamics well. Additionally, spVelo can select biologically significant state driver markers that are validated through enrichment test using oncogenic gene sets from MSigDB [26, 27]. Finally, we present spVelo’s downstream applications, providing new insight into RNA velocity.

## 2. Results

### 2.1 spVelo infers RNA velocity for multi-batch spatial transcriptomics data

spVelo first log-normalizes and smooths the data, and then filters uninformative genes based on their contributions to cell development. Utilizing GO analysis in Extended Data Fig. 2, we demonstrate that the filtered uninformative genes are less enriched for tumor-related pathways (e.g. cytoplasmic translation, structural molecule activity, etc.), compared to other informative genes. spVelo then models unspliced and spliced expression for each gene in a cell as a function of kinetic parameters (transcription, splicing, and degradation rates), latent time, and latent transcriptional state. In each cell, each gene’s latent times are tied via a low-dimension latent variable, following the model assumptions of veloVI [15].

spVelo models the gene expression data with a VAE including two orthogonal encoders. 108 The Multi-Layer Perceptron (MLP) encoder takes the unspliced and spliced expression as input, and outputs the posterior distributions of latent variable. Then, spVelo uses spatial location proximity and distance between batches as the input for a GAT encoder. By adding up the latent space of the two encoders, spVelo is able to jointly model the spatial location and gene expression data. Then by variational posterior inference, spVelo can estimate the kinetic rates and latent time, and then further infer velocity. Additionally, we provide downstream applications including uncertainty quantification, trajectory patterns discovery, state driver markers identification, Gene Regulatory Network (GRN) inference, and temporal cell-cell communication (CCC) inference. A detailed explanation of the spVelo model can be found in the Methods section and the model architecture is shown in Fig. 1. spVelo improves model performance, and provides interpretable results and downstream applications of RNA velocity, suggesting the efficacy of its model design.

**Fig. 1.**
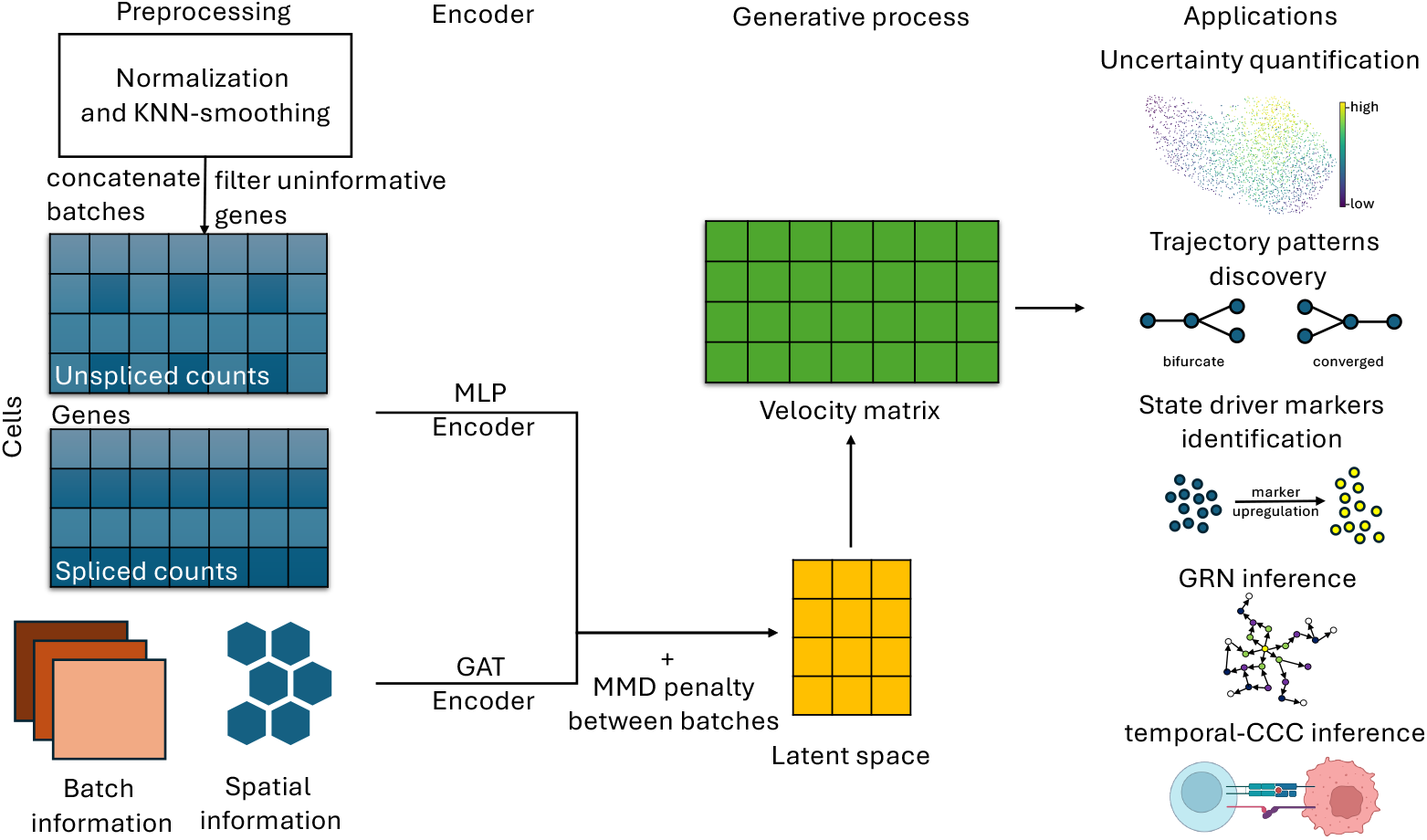
Overview of spVelo. spVelo jointly models the spatial location and gene expression data by using an MLP encoder to encode information from the expression level, and a GAT encoder to encode spatial and batch information. After posterior inference, the velocity matrix can be used for downstream applications.

### 2.2 spVelo infers accurate velocity and trajectory

We first evaluated the performance of spVelo on a spatial dataset simulated from scRNA-seq pancreas data [24] using scCube [28], and a real OSCC dataset [25]. We compared the performance of velocity with other models including stochastic mode and dynamical mode of scVelo [10], veloVI [15], standard mode and annotated mode of LatentVelo [13]. We evaluated the performance of all methods based on the velocity confidence score, transition score, and direction score. The velocity confidence score measures the reliability of inferred velocities, the transition score assesses the probability of true cell-to-cell transition, and the direction score evaluates the consistency of transition directions with known cell type transitions. The three scores are calculated respectively using neighbors of expression data, spatial neighbors in each batch, and mutual nearest neighbors between batches. Detailed explanations of metrics can be found in the Methods section. Since all methods except LatentVelo are restricted to inferring velocity on a per-batch basis, for fairness, we utilized scGen [29] to correct batch effect prior to applying the velocity inference methods. These methods are denoted as scGen + `<method name>` in Fig. 2. For comparing only the per-batch scores (expr scores and spatial scores), we compared spVelo with both scGen-corrected methods and original per-batch methods. Fig. 2 (a) and Fig. 2 (c) show plots of the nine scores for each method by averaging across different seeds and different batches, while Fig. 2 (b) and Fig. 2 (d) show dotplots of only the six per-batch scores for all methods. Here we didn’t compare LatentVelo in the simulated pancreas dataset since it reported errors when input data were in the logcounts format.

**Fig. 2.**
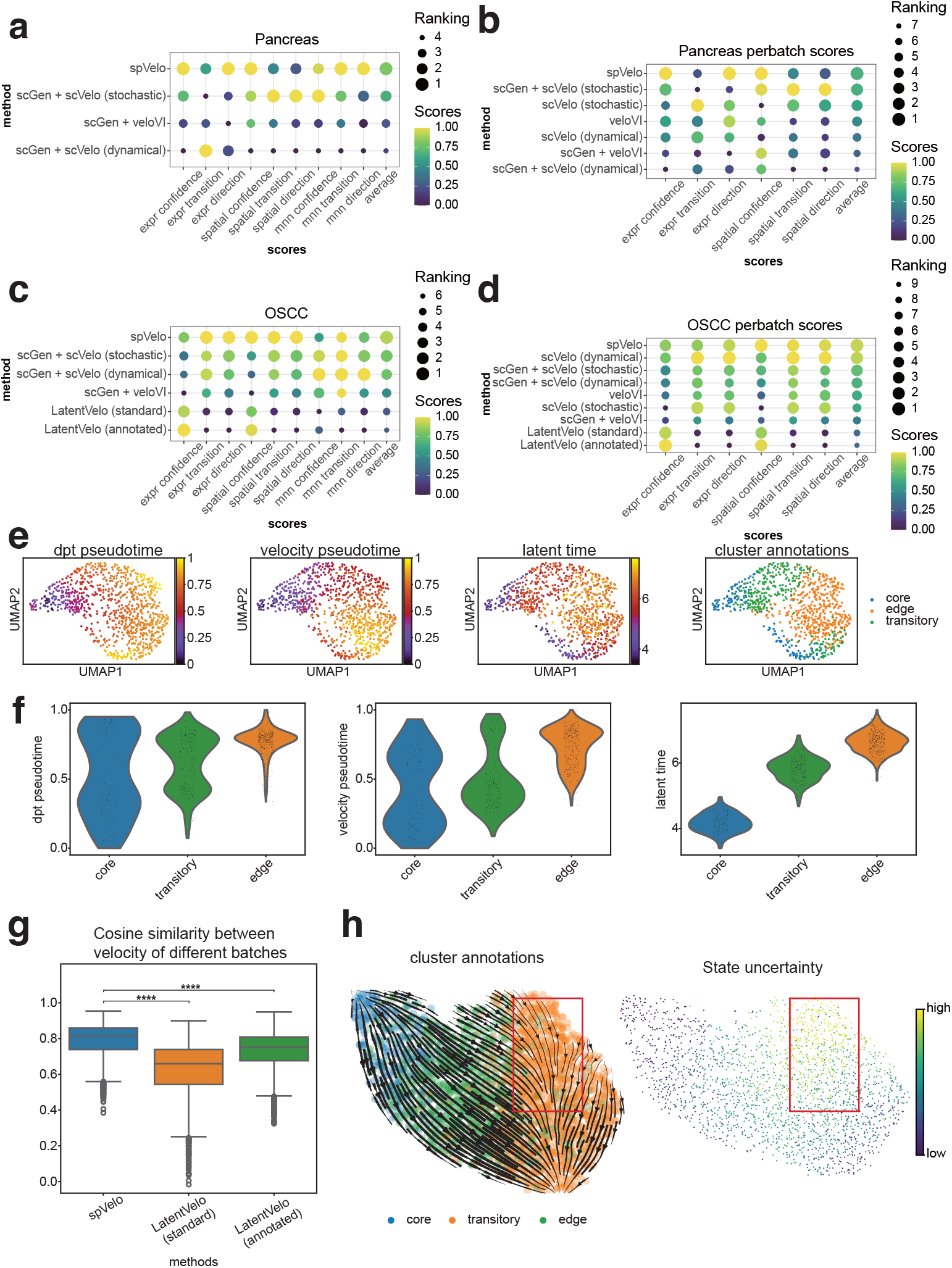
Compare results for simulated pancreas dataset and OSCC dataset. (a) Dotplot of comparing all scores in simulated pancreas dataset. (b) Dotplot of comparing only per-batch scores in simulated pancreas dataset. (c) Dotplot of comparing all scores in the OSCC dataset. (d) Dotplot of comparing only per-batch scores in OSCC dataset. (e) Pseudo-time scatter plot of latent time inferred by spVelo, compared with DPT pseudo-time and velocity pseudo-time. (f) Pseudo-time violin plot of latent time inferred by spVelo, compared with DPT pseudo-time and velocity pseudo-time. (g) Comparison of cosine similarity between the velocity of different batches in MNN graph. (h) Streamline plot of trajectory and scatter plot of quantified uncertainty for sample 9 of OSCC dataset. The red frame in the streamline plot indicates the lineage with high uncertainty cells.

Dotplots in Fig. 2 (a-d) demonstrate that spVelo ranks high when compared to all methods, especially in the direction score, which is the most important score for evaluating velocity’s performance in trajectory inference. Overall, spVelo consistently achieves the highest average scores across all datasets, as illustrated in the final column. This highlights spVelo’s ability to accurately capture the underlying cellular dynamics. All scores are visualized in Extended Data Fig. 4 and Extended Data Fig. 5. We further performed an ablation test to remove spatial information from our model. Results are visualized in Extended Data Fig. 1 and reveal that the integration of spatial information during model training significantly improves the performance of velocity and trajectory inference.

To evaluate the ability of spVelo to correct batch effect in RNA velocity inference, we calculated the cosine similarity between the velocity of mutual nearest neighbor cells in different batches. The comparison result to LatentVelo is visualized in Fig. 2 (g). The boxplot reveals that spVelo infers significantly more coherent velocity than LatentVelo. This shows that, with the MMD penalty between latent space of different batches, spVelo is able to infer more coherent velocity between batches. The coherence in velocity may also facilitate more accurate trajectory inference, since the aligned velocities better reflect the true underlying biological processes rather than noises.

Furthermore, we examined the latent time estimated by spVelo and compared it with pseudo-time inferred using Diffusion Pseudo-Time (DPT) [30] and pseudo-time inferred using diffusion-based random walk on RNA velocity matrix. The results are shown as the scatter plots and violin plots in Fig. 2 (e-f), and all other results are shown in Extended Data Fig. 10. The plots reveal that our inferred latent time is distinct between different cell types and better matches with ground truth.

### 2.3 spVelo quantifies uncertainty for cell state

Since spVelo is a generative model, the distribution of its latent space can be used for uncertainty quantification. Inspired by VeloVAE [31], we calculated differential entropy on the variance of latent space. Since the latent space is a low-dimension representation of cells, the differential entropy can be used as the uncertainty measurement for cell state [32], where higher differential entropy indicates a higher uncertainty score.

We visualized the streamline plot of trajectory and the scatter plot of quantified uncertainty for sample 9 of the OSCC dataset in Fig. 2 (h). Results of other samples are visualized in Extended Data Fig. 6. The plots reveal that some edge cells show higher uncertainty levels. These cells are mostly located at the starting area of the lineage in the red frame, suggesting heterogeneity in the edge cells. This observation also matches with the interpretation in VeloVAE that multi-potent progenitor cells have higher cell state uncertainty [31]. As a result, the uncertainty quantification from spVelo allows researchers to identify and examine the regions with high variability, and further understand intricate biological mechanisms.

### 2.4 spVelo discovers complex trajectory patterns

In this section, we investigated the trajectory inferred using velocity from different methods. From Fig. 3 (a), spVelo inferred a bifurcate trajectory from sample 12 of the OSCC dataset. To validate the inferred bifurcate trajectory, we visualized how spliced expression varies along with latent time inferred by spVelo in scatter plots. Velocity clusters were calculated by using Leiden clustering [33] on the inferred velocity matrix. Expression data and latent time were calculated by averaging the top five markers of edge (1) cells and edge (2) cells. From the visualized scatter plots in Fig. 3 (c), markers of edge (1) are up-regulated in the first lineage (core (1), transitory (1) and edge (1) cells), while markers of edge (2) are up-regulated in the second lineage (core (1), transitory (2) and edge (2) cells). For distinct comparison, we fitted two lines to the two lineages in the first scatter plot. The t-test between the slopes of the two lines shows the statistical significance of the difference between the two lineages, thereby validating the bifurcate trajectory inferred by spVelo.

**Fig. 3.**
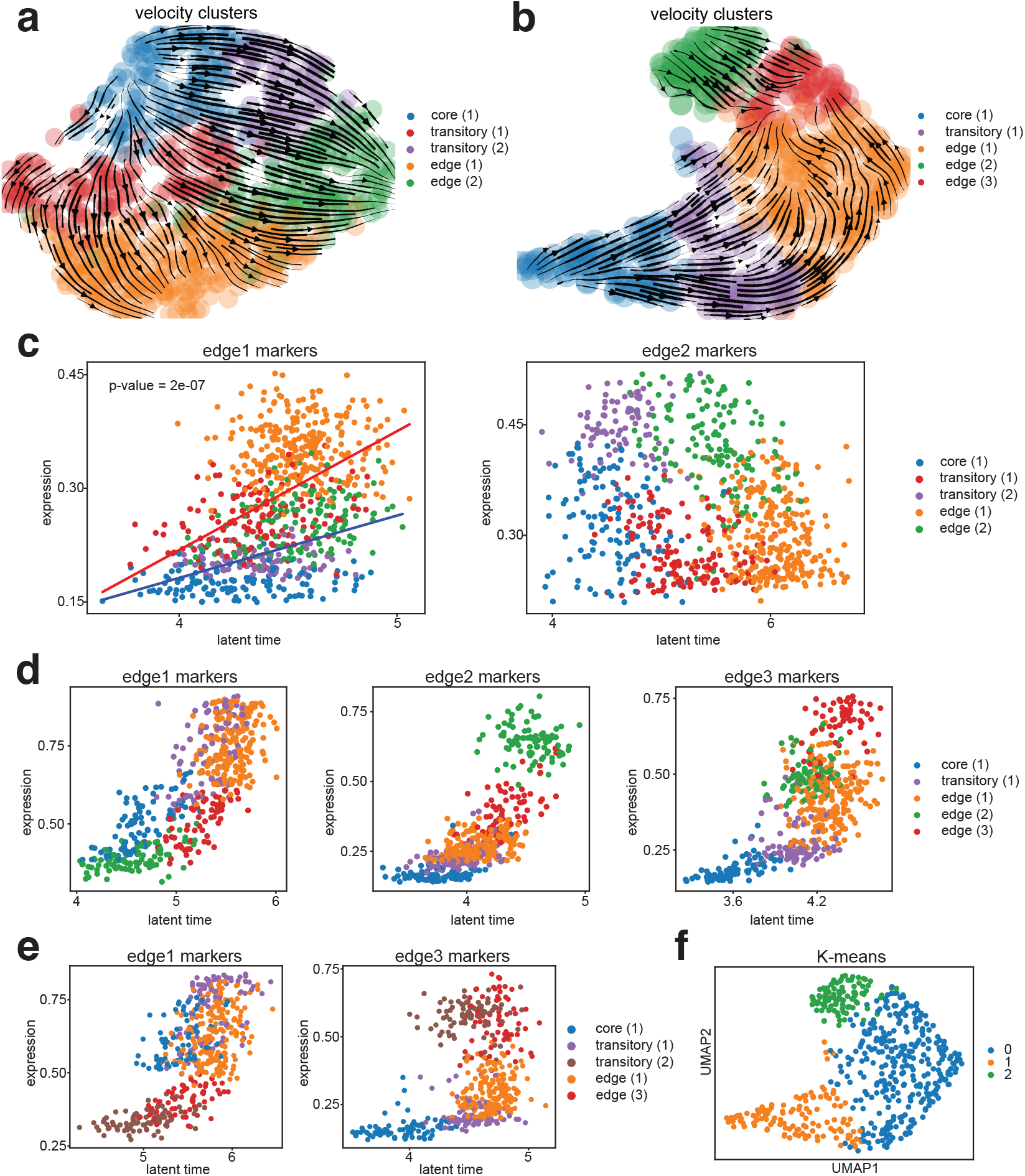
spVelo discovers complex trajectory patterns. (a) UMAP of bifurcate trajectory in sample 12 from the OSCC dataset. (b) UMAP of converged trajectory in sample 4 from OSCC dataset before re-annotation. (c) Scatter plot of how spliced expression varies along with latent time inferred by spVelo in sample 12. Each dot represents a cell, and expression and latent time are calculated by averaging the top five markers of edge (1) cells and edge (2) cells. Linear regression lines are fitted for each lineage in the first scatter plot, with a p value indicating the significance of slope difference. (d) Scatter plot of how spliced expression varies along with latent time inferred by spVelo before re-annotation in sample 4. Each dot represents a cell, and expression and latent time are calculated by averaging the top five markers of edge (1), edge (2), and edge (3) cells. (e) Scatter plot of how spliced expression varies along with latent time inferred by spVelo after re-annotation in sample 4. Each dot represents a cell, and expression and latent time are calculated by averaging the updated top five markers of edge (1) and edge (3) cells. (f) UMAP of K-means clustering.

Additionally, for sample 4 of the OSCC dataset, spVelo inferred a converged trajectory as shown in Fig. 3 (b). The clustered results indicated three edge sub-types. Similarly, we visualized the scatter plots of averaged spliced expression and latent time in Fig. 3 (d). However, upon closer examination, the expression patterns of edge (2) are more consistent with transitory (2) cells, since they transition into edge (3). As a result, we re-annotated edge (2) into transitory (2) and presented the scatter plots after re-annotation in the lower half of Fig. 3 (e).

In the left panel of Fig. 3 (e), the first lineage (core (1), transitory (1), and edge (1) cells) expresses edge1 markers at a higher level, while the second lineage (transitory (2) and edge (3) cells) expresses at a lower level. The right panel of Fig. 3 (e) shows the opposite for edge3 markers. We further performed K-means clustering with the concatenate of latent time matrix and gene expression matrix as input and n_clusters set as 3. From the visualization in Fig. 3 (f), previous edge (2) cells should be separated from edge (3) cells. As a result, this updated information aligns the cell classifications with expression dynamics and more accurately reflects the cell type transitions, further supporting spVelo’s capability in identifying complex cellular dynamics and refining cell type classifications. The trajectory plots of all other samples on UMAP embedding and spatial coordinates are visualized in Extended Data Fig. 6-8.

### 2.5 spVelo improves genes’ fit and selects biologically important state driver markers

Multiple rate kinetics (MURK) genes are defined as genes with transcriptional boosts [20]. Their expression levels increase rapidly during specific cellular states. Models with simple assumptions may fail to capture their complex dynamics. These up-regulating boosts would lead to down-regulation estimations, and may further lead to reversed estimations of cellular transitions [11]. Possible solutions include manually removing the MURK genes that violate model assumption [20]. However, this removal risks the loss of biologically informative genes that are crucial for velocity and trajectory inference.

To address this limitation, we evaluated the capacity of spVelo in inferring the kinetic rates of MURK genes. In Fig. 4 (a), we visualized phase portraits of five MURK genes from the OSCC dataset, showing the robustness of spVelo in capturing the non-linear dynamics and estimating complex kinetics. By fitting the MURK genes, spVelo provides a more accurate representation of the underlying biological process. We also visualized phase portraits of state driver markers selected from the simulated pancreas dataset in Fig. 4 (b). This further demonstrates spVelo’s ability to accurately fit genes’ dynamics.

**Fig. 4.**
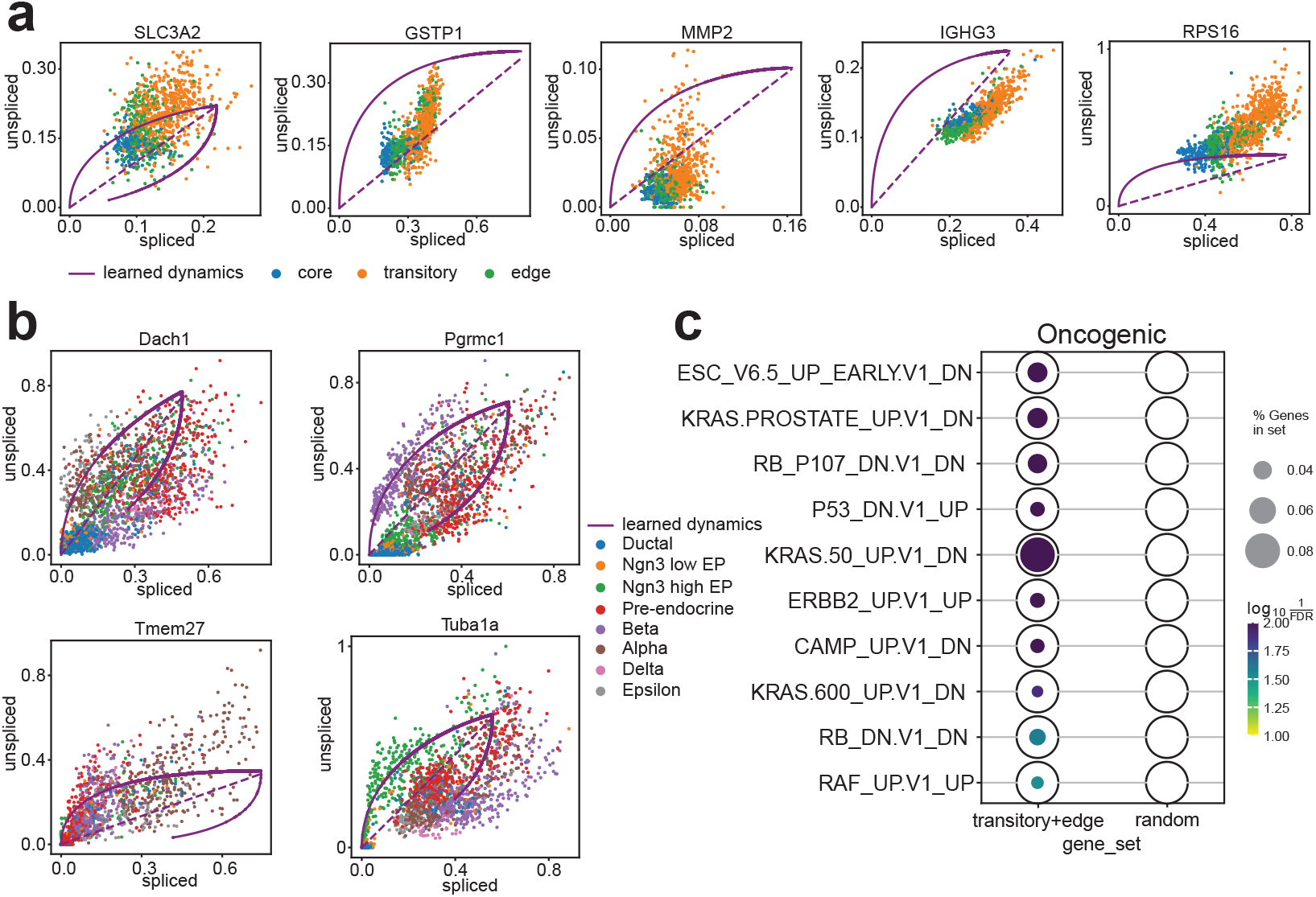
spVelo fits genes’ dynamics well. (a) Phase portraits of five MURK genes from the OSCC dataset. (b) Phase portraits of state driver markers selected from simulated pancreas dataset. (c) spVelo selects biologically significant state driver markers, verified by gene set enrichment analysis using MSigDB.

Furthermore, we examined the biological significance of state driver markers selected by spVelo. Based on the velocity estimation, state driver markers are defined as genes pivotal in driving cellular state transitions. Here we utilized a t-test on the estimated velocity matrix to select state driver markers and used oncogenic gene sets from MsigDB [26, 27] for gene set enrichment analysis (GSEA). We visualized the GSEA results through a dotplot in Fig. 4 (c). The first column of the dotplot is state-driver markers selected from transitory and edge cells, and the second column is the same number of randomly selected genes from the dataset, serving as a control group. The dotplot demonstrates that the state driver markers are significantly enriched in oncogenic pathways compared to the random gene set, proving spVelo’s ability to select state driver markers that play a crucial role in cancer progression. These state driver markers can potentially serve as targets for therapeutic intervention.

### 2.6 spVelo infers gene regulatory networks by in-silico gene deletion

Gene regulatory network (GRN) inference is a popular area since it is critical for understanding transcription. Here we present spVelo’s downstream application in GRN inference. Inspired by [34], we employed an in-silico gene deletion approach. We inferred the velocity before and after removing *EGFR*, a gene known for prompting OSCC cell proliferation, metastasis, invasion, and apoptosis resistance [25, 35, 36]. To quantify the impact of *EGFR* deletion, we calculated the gene-wise cosine similarity between the two velocity matrices obtained before and after in-silico perturbation. The comparison between EGFR target genes and target genes of other genes is visualized in Fig. 5 (a). The boxplot reveals that direct EGFR targets (defined by the transcription factor target gene sets from MsigDB [26, 27]) are more impacted by the in-silico deletion of *EGFR* compared to other target genes. The results suggest that with in-silico perturbation, spVelo may identify regulatory relationships and enable the identification of critical genes driving biological processes, thus contributing to understanding the mechanisms underlying disease progression.

**Fig. 5.**
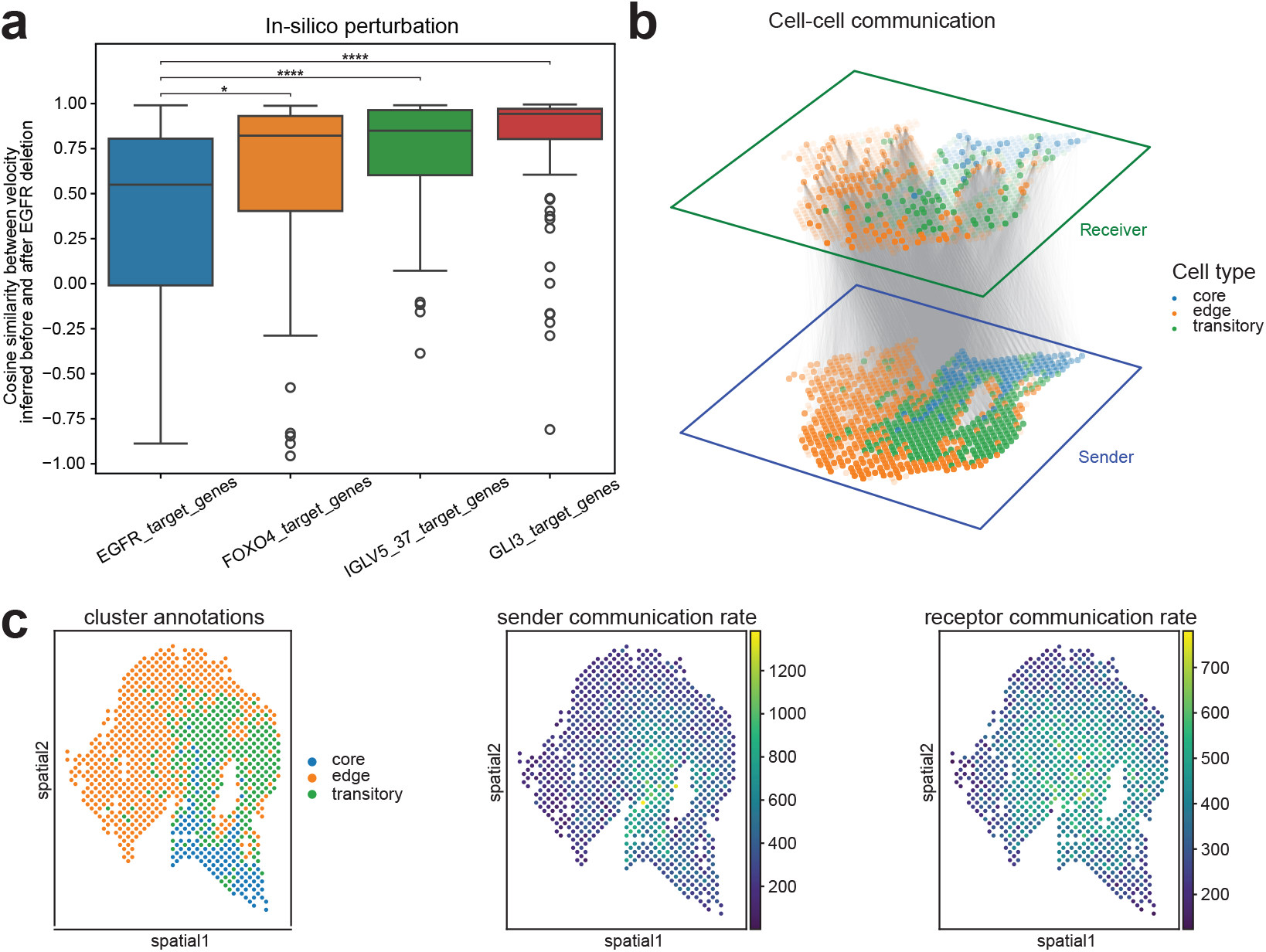
Downstream applications of spVelo. (a) The Y-axis is the cosine similarity calculated of each gene’s velocity before and after in-silico perturbation. The boxplot compares the cosine similarity between EGFR target genes and other target genes. (b) 3D plot of inferred cell-cell communication. (c) Temporal cell-cell communication inferred with velocity from spVelo. From left to right: spatial scatter plot of sample 2 from OSCC, scatter plot with sender communication rate, scatter plot with receptor communication rate.

### 2.7 spVelo enables temporal cell-cell communication inference

Inspired by CytoSignal and VeloCytoSignal [37], we inferred cell-cell communication (CCC) and temporal CCC using spVelo. Detailed steps of CCC inference can be found in the Methods section. Here we used the ligand-receptor gene pair (*ANXA1, EGFR*) for CCC inference. The inferred spot-level CCC is visualized in Fig. 5 (b), where lines between sender and receptor cells indicate communications between them. From Fig. 5 (b), few core cells are receptors, consistent with the cell transition ground truth provided by [25].

Given the significance of CCC in dynamical processes, we quantified spatial-temporal changes in signaling activities to understand the role of CCC in cell state transition.

Utilizing RNA velocity, we inferred temporal CCC and visualized the sender and receptor communication rate in Fig. 5 (c), and the other results are shown in Extended Data Fig. 11. Fig. 5 (c) reveals that sender communication rates are higher in core and transitory cells, while receptor communication rates are higher in transitory and edge cells. This result also aligns with ground truth, demonstrating that spVelo effectively captures temporal dynamics in cell-cell communications. This helps elucidate the signaling networks in both static and developmental contexts, enabling researchers to better understand the timing of critical cellular interactions.

## 3 Discussion

RNA velocity has emerged as a new approach for inferring cellular trajectory and understanding dynamical processes. Meanwhile, spatially resolved transcriptomics combines gene expression with spatial context, offering insights into cellular architectures. However, existing RNA velocity methods fail to utilize these spatial insights, particularly in large-scale, multi-batch datasets. Here, we introduce spVelo, a novel RNA velocity inference method for multi-batch spatial transcriptomics datasets. Our extensive analysis proves its accuracy and interpretability in velocity and trajectory inference.

Existing methods exhibit several limitations when applied to large-scale spatial datasets. All methods are developed for scRNA-seq and are unable to utilize the spatial information. Among the compared methods, scVelo suffers from strict assumptions and simple modeling, making it unable to capture complex dynamics. This results in oversimplified or inaccurate trajectory inference. On the other hand, veloVI presents a complex VAE-based model with a time-dependent transcriptional rate. However, it fails to infer RNA velocity from multi-batch datasets. LatentVelo is scalable to multi-batch datasets by incorporating batch information into its model, yet fails to infer coherent velocity between batches and infers inaccurate trajectory.

spVelo overcomes the above limitations. With its design of combining VAE with GAT, spVelo is capable of leveraging the information from both spatial location and expression data. Additionally, by introducing an MMD penalty between batches, spVelo is able to infer coherent velocity from multi-batch datasets. Consequently, spVelo more accurately infers velocity and trajectory from large-scale datasets, effectively capturing the underlying dynamics of tissues.

We further provided downstream applications utilizing the velocity inferred by spVelo. Firstly, we demonstrated that the generative modeling of spVelo enables interpretable uncertainty quantification. Secondly, we discovered complex trajectory patterns and further discovered possible cell type refinement. Thirdly, we selected state driver markers and proved their biological significance. Fourthly, we inferred the Gene Regulatory Network utilizing an in-silico gene deletion approach. Finally, we inferred temporal cell-cell communications that are consistent with ground truth. Therefore, RNA velocity inferred by spVelo offers new biological insight into cellular dynamics and exhibits great promise for future explorations.

## 4 Methods

### 4.2 Problem definition

In the RNA velocity inference problem, we denote the spliced expression matrix as *S*^*N×G*^ and unspliced expression matrix as *U* ^*N×G*^, where *N* represents the number of cells and *G* represents the number of genes. We use *X*^*N×*2^ to represent the spatial locations of the cells. With these as input, spVelo aims to learn a model *M*, which can infer the cell-by-gene velocity matrix as: *V* ^*N×G*^ = *M* (*S, U, X*). The model can simultaneously infer cell-gene-specific latent time *t*_*ng*_, transcriptional state *k*, and kinetic rates including gene-state-specific transcription rate *α*_*gk*_, gene-specific splicing rate *β*_*g*_, and gene-specific degradation rate *γ*_*g*_. Here transcriptional state *k* ∈{1, 2, 3, 4}, where *k* = 1 indicates induction, *k* = 2 indicates the induction steady state, *k* = 3 indicates repression, and *k* = 4 indicates the repression steady state.

### 4.2 spVelo model specification

Following [10] and [15], spVelo assumes that for each gene, cells first go through an induction state where spliced and unspliced expression increases. Then cells reach an induction steady state, and then at a switching time, the system switches to a repression state where spliced and unspliced expression decreases. Finally, cells reach a repression steady state with no expression.

By solving the ordinary differential equations [10], the estimated unspliced and spliced abundance at time *t*_*ng*_ for cell *n* and gene *g* is defined as:

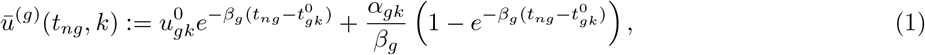

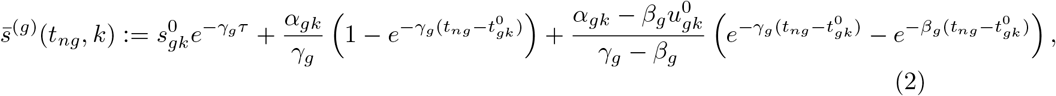

where 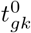 denotes the initial time of the system in state 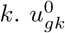 and 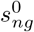 denotes the estimated initial unspliced and spliced expression of gene *g* in state *k*, i.e. 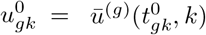 and 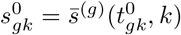.

Transcription rate *α* is assumed to be time-dependent with parameters *α*_0_, *α*_1_, *λ*_*α*_:

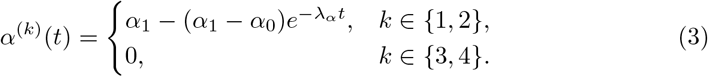

For future conciseness, we still write the gene-state-specific transcription rate 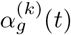 as *α*_*gk*_.

For *k* = 1 (induction state), we have 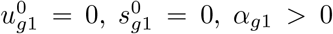 and 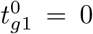 by definition. Thus (6) and (7) can be simplified into

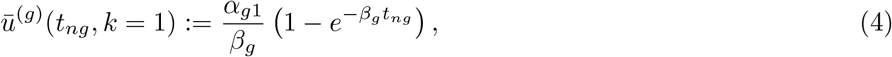

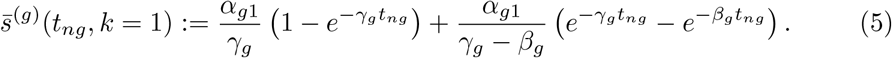

For *k* = 2 (induction steady state), the unspliced and spliced expression is defined as the limit of the induction state as time approaches ∞:

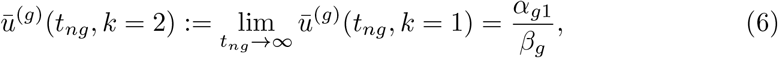

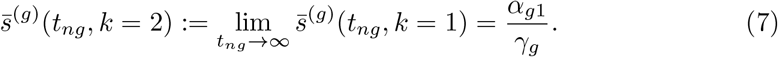

For *k* = 3 (repression state), we have *α*_*g*3_ = 0 and 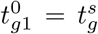,where 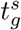 is the gene-specific switching time from the induction phase to the repression phase. Thus (6) and (7) can be expressed as

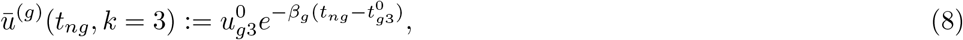

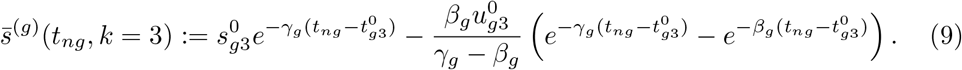

Similarly, *k* = 4 (repression steady state) is defined as the limit of the repression state, resulting in

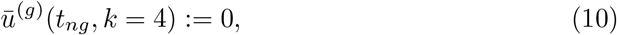

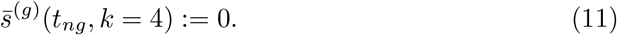

### 4.3 spVelo generative process

The generative modeling of spVelo combines a Variational Autoencoder (VAE) [21] inspired by [15], with a Graph Attention Network (GAT) [22]. We assume the following generative process to model the underlying dynamics of the unspliced expression *u*_*ng*_ and spliced expression *s*_*ng*_.

For each cell *n* and gene *g*, we use a low-dimension latent variable *z*_*n*_ to summarize the latent state of each cell (default *d* = 10). *z*_*n*_ is the sum of latent space from VAE and GAT, modeling both expression data and spatial location. Let

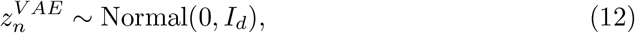

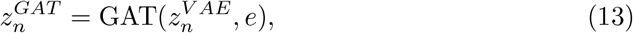

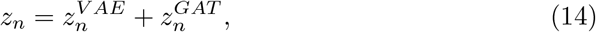

where *e* denotes the edges input to GAT. In GAT modeling, 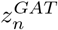 is constructed based on a graph structure where edges represent relationships between cells. The edges are composed of two parts: The first part of the edges is calculated using k Nearest Neighbors (kNN) on the spatial coordinates. We compute the edges in each batch and concatenate across all batches. The second part of edges is calculated across different batches using Mutual Nearest Neighbors (MNN) on the expression data. The distance of MNN is defined as the optimal transport (OT) matrix, quantifying the correspondence between samples in different batches [38]. The metric cost matrix in the OT problem is calculated as the Euclidean distance between batches. By combining the two parts of edges, the GAT module effectively captures spatial information together with relationships between batches. The number of neighbors for both parts is default set as 15. The effects of hyper-parameter tuning are visualized in Extended Data Fig. 9.

We then use a Dirichlet distribution to model state assignment probability *π*_*ng*_. The settings are based on VeloVI. The state *k*_*ng*_ is then defined as the state with the highest state assignment probability.

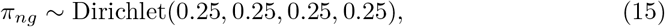

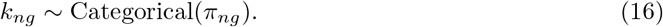

Latent time *t*_*ng*_ is modeled as a state-specific function of latent state *z*_*n*_:

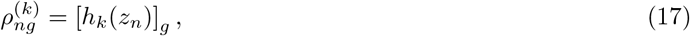

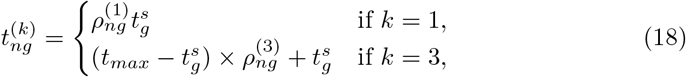

where *t*_*max*_ := 20 fixes the time scale across genes. *h*_*k*_ : ℝ^*d*^→ (0, 1)^*G*^ is parameterized as a state-specific fully connected neural network.

Finally, we assume the observed expression data are sampled from Normal distributions as

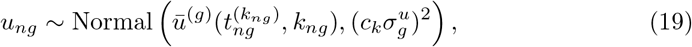

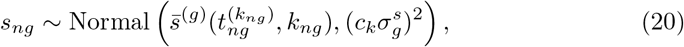

where *c*_*k*_ is a state-dependent scaling factor on the variance. As default, *c*_*k*_ = 1 for *k* = 1, 2, 3 except for *c*_4_ = 0.1 in the repression steady state.

### 4.4 spVelo posterior inference

#### Variational posterior

Let *θ* be the set of parameters including kinetic rates (*α, β, γ*), switching time *t*^*s*^, and neural network parameters. We use variational inference [21] to approximate the posterior distribution. The posterior distribution is posited as

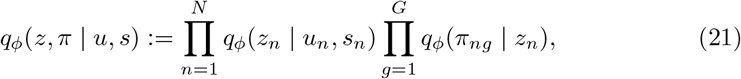

where dependencies are specified using neural networks with parameter set *ϕ*.

Integrating over choice of transcriptional state *k*_*ng*_, the likelihoods for spliced and unspliced transcript abundances are Gaussian mixture models:

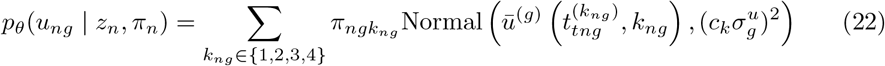

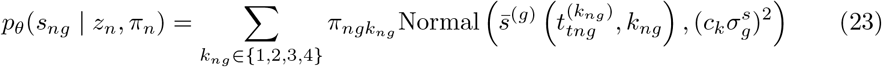

#### Optimization

The objective function is composed of three terms

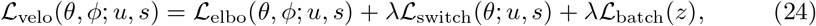

where ℒ_elbo_ is the negative evidence lower bound [39] of *logp*_*θ*_(*u, s*), ℒ_switch_ is a penalty that regularizes the location of transcriptional switch in the phase portrait, and ℒ_batch_ is an MMD penalty that regularizes the latent space between different batches. As default, the penalty weight *λ* = 2. In more detail, we denote *b*_1_, *b*_2_ as a pair of different batch IDs, *z*_*b*_ as the latent space of batch *b*, and *u*^*^ and *s*^*^ as the median unspliced and spliced expression for each gene,

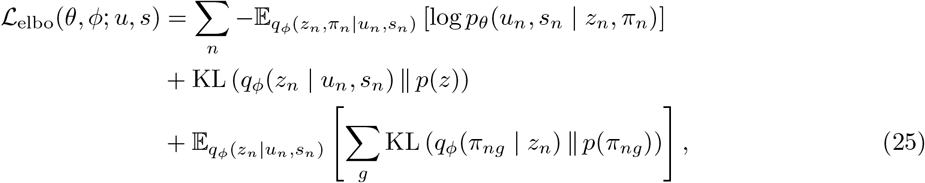

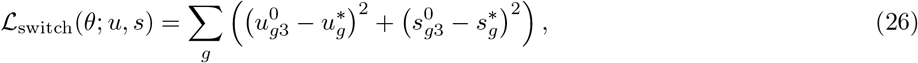

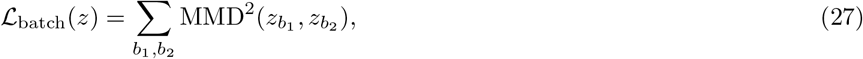

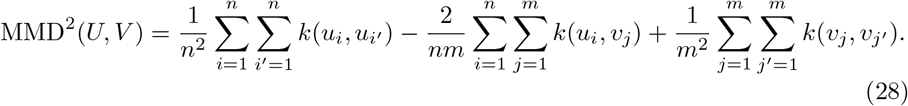

Here *k*(*x, y*) denotes a Gaussian kernel, i.e., 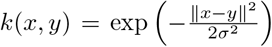,where *σ* is a bandwidth parameter and ∥*x* − *y*∥ is the Euclidean distance between *x* and *y*.

To optimize ℒ_velo_, we use stochastic gradients [21] and Adam optimizer with weight decay [40]. We set the number of epochs as 2,000.

#### Velocity inference

After fitting the parameters, the cell-gene-specific state assignment is calculated as the posterior mean:

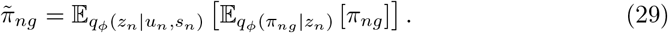

The cell-gene-specific latent time is calculated as

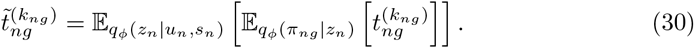

RNA velocity is calculated as a function of the variational posterior

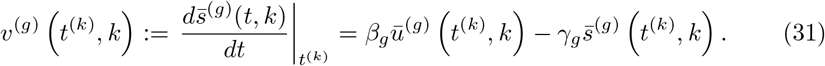

### 4.5 Uncertainty quantification

Uncertainty of latent state is calculated as the differential entropy of latent space:

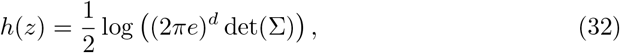

where *d* is the dimension (default as 10) and Σ is the variance matrix of the latent space.

### 4.6 Temporal cell-cell communication inference

The spatial interaction score is defined as the co-expression of ligand and receptor genes within close spatial proximity. Here we select a ligand-receptor gene pair from OmniPath [41] and denote spliced expression matrix as *S*, and denote a pair of ligand and receptor genes as *l* and *r*.

For cells *i* and *j*, we calculate the LRscore as:

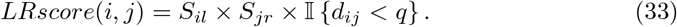

For cell types *A* and *B*, we calculate the LRscore as:

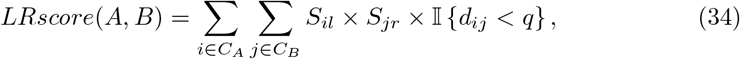

where *C*_*A*_ refers to all cells in cell type A, and *S*_*il*_ refers to the expression value of gene *l* in cell *i*. In the indicator function, *d*_*ij*_ refers to the Euclidean distance between the spatial location of cell *i* and cell *j*, and *q* refers to a user-defined threshold, default as 30. After calculating scores between cell types, we randomly permuted cell types 50 times and performed False Discovery Rate (FDR) correction.

The spatial-temporal interaction score is defined as the time derivative of LRscore and calculated as follows:

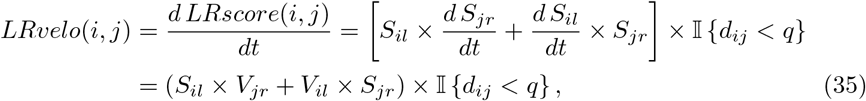

where *V* refers to the inferred velocity matrix.

### 4.7 Metrics explanations

To evaluate the performance of inferred velocity, we calculated three different types of scores, inspired by VeloAE [42]. For each pair of cell types (*A, B*), the scores are calculated for the boundary scores, referring to cells of cell type A with cell type B in the neighborhood, i.e., *C*_*A*→*B*_ ={ *c* ∈ *C*_*A*_ |∃*c*^*′*^∈ *C*_*B*_ ∩ *N* (*c*)}. Here *C*_*A*_ denotes all the cells of cell type *A* and *N* (*c*) denotes neighbor cells of *c*.

#### 1. Confidence score

Confidence score for cell *c* from cell type *A* with regards to cell type *B* is defined as

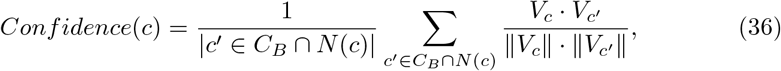

where *V*_*c*_ is the velocity vector of cell *c*. This is calculated using `scv.tl.velocity_confidence`. Then the confidence score for cell type *A* is calculated as the average of *Confidence*(*c*) for all *c* ∈ *C*_*A*→*B*_. It summarizes the consistency of the inferred velocity and a higher confidence score represents better consistency.

#### 2. Transition score

Transition score for cell *c* from cell type *A* with regards to cell type *B* is defined as

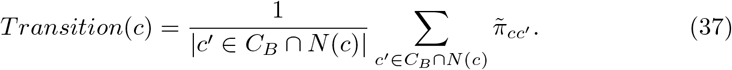

Here 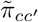 denotes the cell-to-cell transition probabilities calculated from the velocity graph *π*_*cc*_^*′*^ with row-normalization *z*_*c*_ and kernel width *σ*. This is calculated using `scv.tl.velocity_graph and scv.utils.get_transition_matrix`.

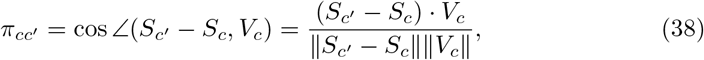

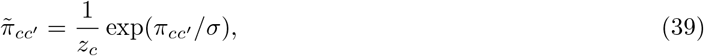

where *S*_*c*_ refers to spliced gene expression of cell *c*. Transition score for cell type *A* is calculated as the average of *Transition*(*c*) for all *c* ∈ *C*_*A*→*B*_, measuring how well the corresponding change in gene expression matches the predicted change. A higher transition score represents a better match.

#### 3. Direction score

Direction score for cell *c* from cell type *A* with regards to cell type *B* is defined as

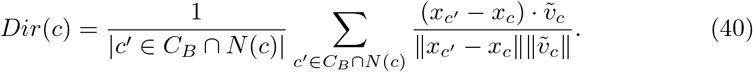

Here *x*_*c*_ and *x*_*c*_^*′*^ are vectors representing cells *c* and *c*^*′*^ in a low-dimension Principal Component Analysis (PCA) space via [43] (number of principal components default as 30). *x*_*c*_^*′*^ − *x*_*c*_ is the displacement in this space, and 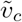 is the projection of velocity into PCA space, calculated using `scv.tl.velocity_embedding`. Denoting 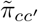 as the transition probability matrix, we have

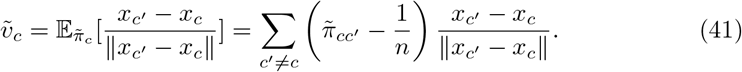

Direction score for cell type *A* is calculated as the average of *Dir*(*c*) for all *c* ∈ *C*_*A*→*B*_, measuring how well the corresponding change in PCA embedding matches the predicted change. A higher direction score represents a better match.

With ground truth cell type transition information as input, the confidence scores are calculated as the average score of all correct cell type transition pairs, while transition scores and direction scores are calculated by averaging scores of correct cell type transition pairs while incorporating a penalty for incorrect transitions by using their negated scores. More discussions of the metrics can be found in Appendix 2.

From the equations, the three scores are all calculated based on local neighborhoods. We compute the kNN graph, spatial graph and MNN graph respectively, incorporating different information in model comparison. As default, the neighbor size is set as 30.

Inspired by LatentVelo [13], we also measure the cosine similarity of MNN cells in different batches to evaluate batch effect correction of RNA velocity. Let *C*_*b*_ be all the cells in batch *b* and *N* ^*MNN*^ (*c*) be MNN of cell *c*, the velocity coherence score for cell *c* is defined as:

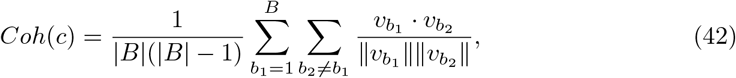

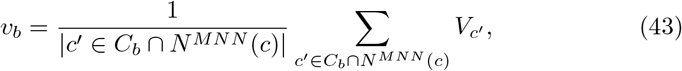

where *B* denotes the set of batches in the dataset and (*b*_1_, *b*_2_) denotes a pair of different batch IDs. Then the final velocity coherence score is calculated as the average of 100 randomly selected cells.

### 4.8 Baseline model explanations

In the model comparison process, we consider eight baseline methods (settings) in total for comparison, including standard and annotated mode of LatentVelo, stochastic and dynamical mode of scVelo, veloVI, and scGen-corrected scVelo and veloVI. The order of these methods (settings) is random.

LatentVelo [13] uses a VAE that embeds unspliced and spliced abundances of RNA into latent space and dynamics on the latent space are described as a neural ODE. By learning a shared latent space for multiple batches, LatentVelo enables batch effect correction from a dynamic view. The annotated mode of LatentVelo incorporates cell type information by modifying the prior.

The stochastic mode of scVelo [10] treats transcription, splicing, and degradation as probabilistic events and approximates the Markov process using moment equations. By using both first- and second-order moments, scVelo (stochastic) can utilize both relationships and covariation between unspliced and spliced mRNA abundances. The dynamical mode of scVelo solves the ODEs with a likelihood-based expectation-maximization framework, iteratively estimating the parameters of kinetics rates, transcriptional state and cell-internal latent time.

veloVI [15] treats unspliced and spliced abundances of RNA for each gene as a function of kinetic parameters, latent time, and latent transcriptional state. It further treats latent time as tied via a low-dimension latent variable. veloVI uses a VAE architecture and outputs posterior distribution over estimated velocity.

For batch effect correction settings, since current RNA velocity methods require cell-by-gene spliced and unspliced counts as input, only batch effect correction methods that return a corrected and reconstructed gene matrix can be used. As a result, we used scGen [29] for batch effect correction, as recommended by scIB [44].

In the scGen-corrected models, we followed the approach taken by [13, 45]. Since we need to simultaneously correct spliced and unspliced counts, we perform batch effect correction on the sum of these counts. Denote spliced and unspliced count as *S* and *U*, we define the sum matrix as *M* = *S* + *U*, and ratio matrix as 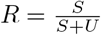.scGen batch effect correction is performed on log-normalized *M* with the default settings, and we get the corrected matrix 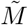.To recover corrected spliced and unspliced expression, we multiply 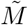 with *R* or 1 − *R*.

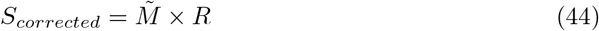

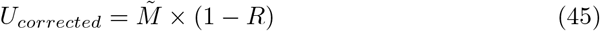

Then RNA velocity is estimated as before.

### 4.9 Experiment design

For the simulated dataset, we followed the tutorial from scCube [28] and generated random spatial patterns for cell types with a reference-free strategy. Extra analysis of data simulation can be found in Appendix 1 and Extended Data Fig. 3. We used the scRNA-seq pancreas dataset [24] for this simulation. For real OSCC dataset, we filtered all noncancer (nc) cells, following the preprocessing step in [25].

For both simulated pancreas dataset and real OSCC dataset, we followed the preprocessing guidelines from scVelo [10]. We normalized the count matrices to the median of total molecules across cells and filtered genes with less than 20 expressed counts commonly for spliced and unspliced mRNA, followed by log-transforming the data and selecting the top 2,000 highly variable genes. Then we calculated a nearest neighbor graph (with 30 neighbors) based on Euclidean distances in principal component analysis space (with 30 principal components) on spliced logcounts. We computed first- and second-order moments (means and uncentered variances) for each cell across its 30 nearest neighbors.

Following [15], we min-max scaled the unspliced and spliced expression to the unit interval and applied the steady-state scVelo model. Finally, we filtered the genes with negative steady-state ratio and *R*^2^ statistic below a user-defined threshold (default as 0.2). Then the remaining genes are used for velocity inference.

In model comparison, we followed the tutorials of all methods.

## 5 Data availability

We summarize the sources and statistics of all datasets we used in Supplementary File 1. All the public datasets can be accessed based on the links in this file.

## 6 Reproductivity and Code availability

We relied on Yale High-performance Computing Center (YCRC) and utilized one NVIDIA A5000 GPU with up to 30 GB RAM for model training. The codes of spVelo can be found at https://github.com/VivLon/spVelo. We follow the MIT license for usage.

7 Acknowledgements

We thank Hanshu Yu for providing suggestions from a biological view.

## 8 Author contributions

T.L. and W.L. designed this study. W.L. and T.L. designed the model. W.L. ran all the experiments. W.L., T.L., L.X., and H.Z. wrote the manuscript. L.X. and H.Z. supervised this work.

