## Appendix for "spVelo: RNA velocity inference for multi-batch spatial transcriptomics data"

### 1 Extra analysis of data simulation

Other than scCube [1], we also considered scDesign3 [2] for simulating spatial datasets. In our implementation, we used sample 2 of the OSCC dataset [3] and followed the step-by-step tutorial of scDesign3 to simulate a new spatial dataset. However, the simulated dataset exhibits unreasonable expression data, as shown in the scatter plots in Extended Data Fig. 3. The relationship between spliced expression levels and unspliced expression levels is incorrect.

### 2 Discussion of metrics

While the transition score and direction score may seem similar, they measure different aspects and serve as a complement to each other. The transition score is a direct assessment of how well the inferred velocity predicts changes in gene activity. However, the gene expression space is often high-dimensional and noisy. The results might be sensitive to noise or irrelevant features. On the other hand, the direction score measures the cosine similarity between the inferred velocity and the change in a low-dimension PCA space. This focuses on more general trends, yet is unable to reflect subtle but important biological information. As a result, using the two scores simultaneously can provide a more complete picture of how well the inferred velocity aligns with true biological processes and offers complementary insights into the performance of velocity inference.

In the dimension reduction process of direction score calculation, we used Principal Component Analysis (PCA) [4] instead of Uniform Manifold Approximation and Projection (UMAP) [5] that was used in [6, 7]. The reason for our choice is: Firstly, PCA has a higher dimension than UMAP, thus preserving more information from the original dataset. Secondly, PCA is deterministic while UMAP is stochastic, so PCA can provide more consistent results. To this end, we chose PCA over UMAP for direction score calculation.

#### 3 Supplementary figures

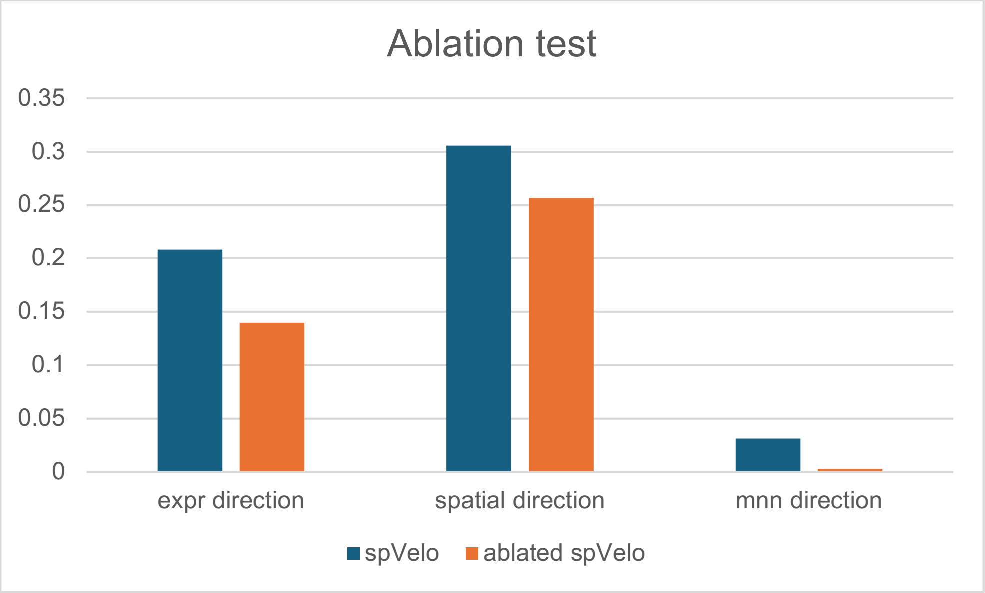


**Extended Data Fig. 1** Ablation test of spVelo in OSCC dataset, comparing expression direction, spatial expression, and mnn direction score between spVelo and spVelo without inputting spatial information.

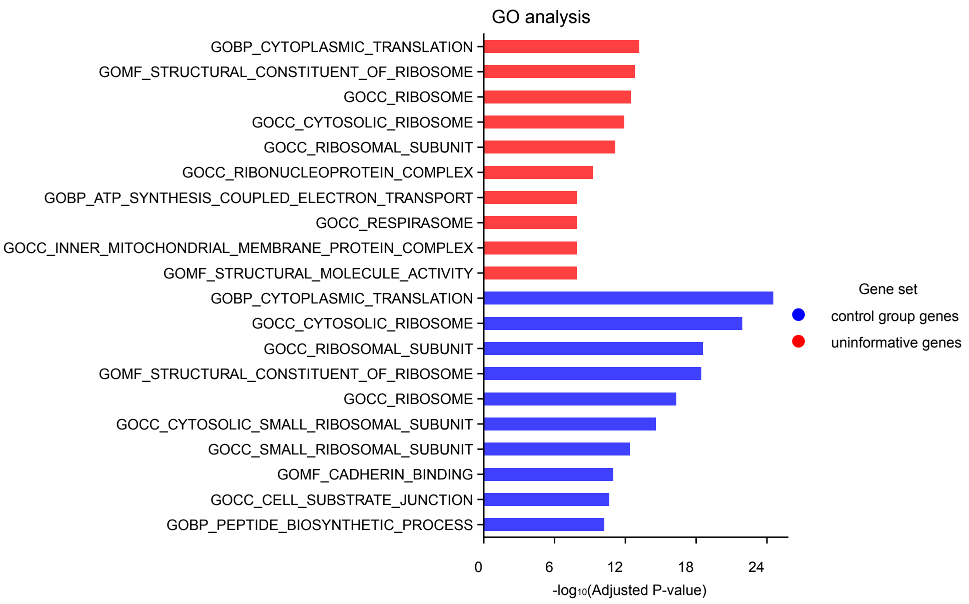


**Extended Data Fig. 2** Dotplot of GO analysis for comparing filtered uninformative genes with control group genes. The control group is randomly selected from informative genes, with the same number of genes as in the uninformative genes.

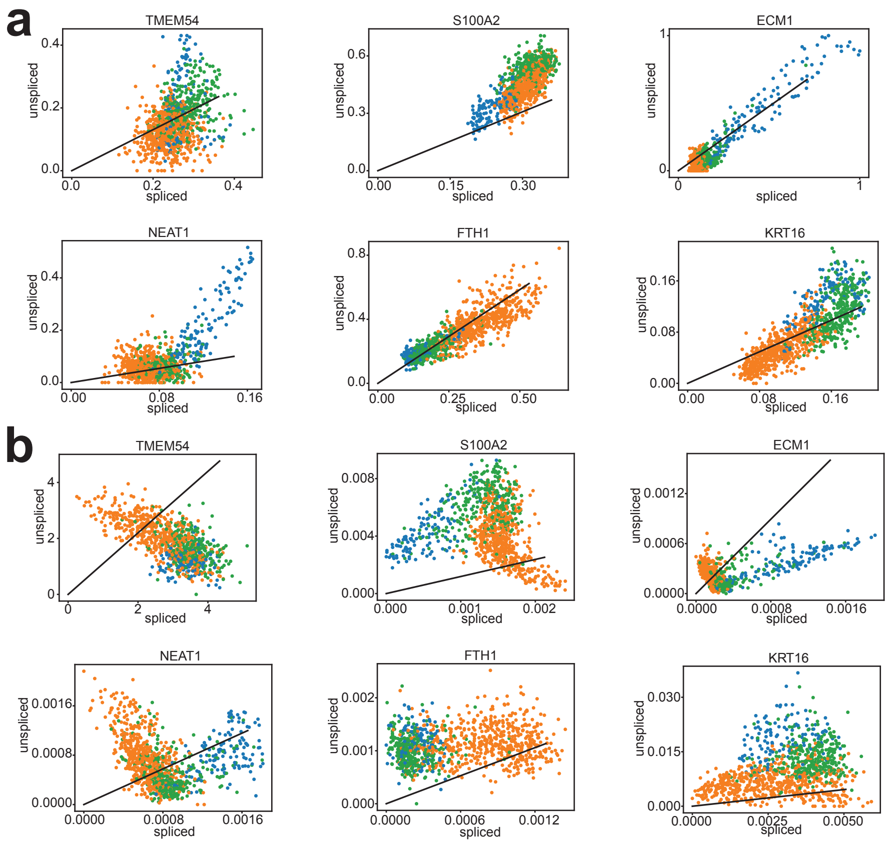


**Extended Data Fig. 3** scDesign3 simulation data quality. (a) Scatter plot of genes from the original dataset. (b) Scatter plot of genes from scDesign3 simulated dataset.

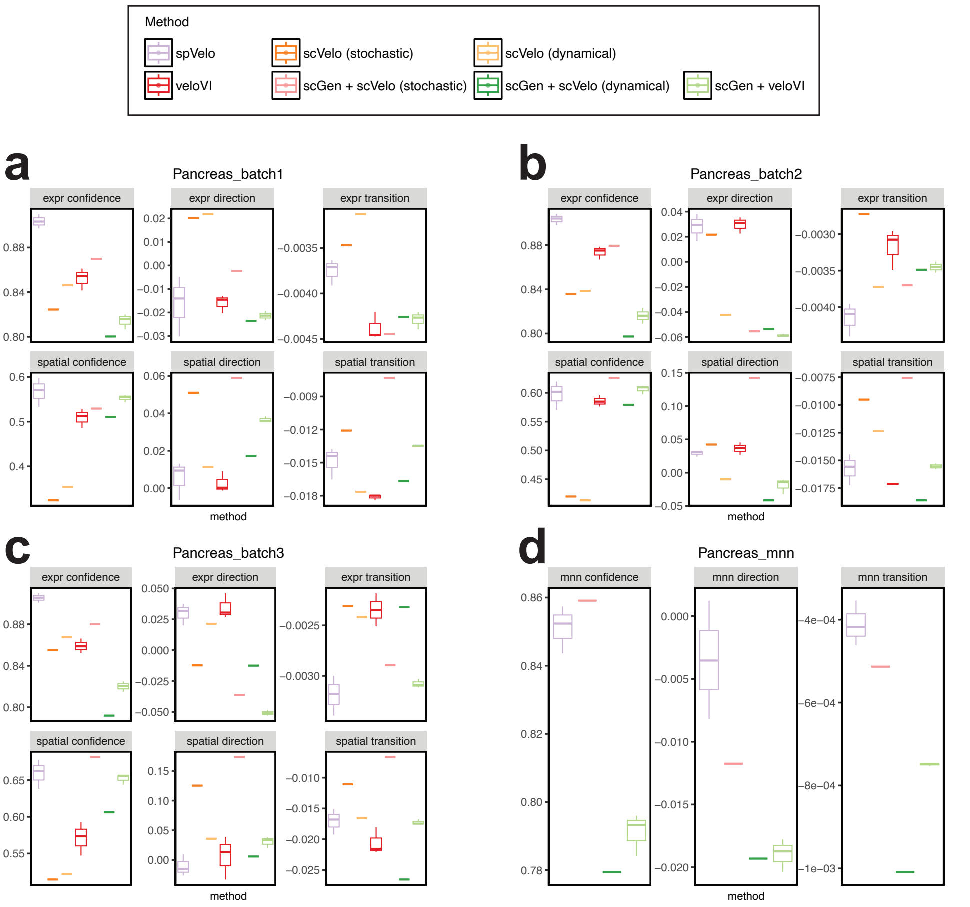


**Extended Data Fig. 4** Boxplots of all comparing scores in simulated pancreas dataset. (a-c) Perbatch scores from batch 1 to batch 3 of simulated pancreas dataset. (d) MNN scores of simulated pancreas dataset.

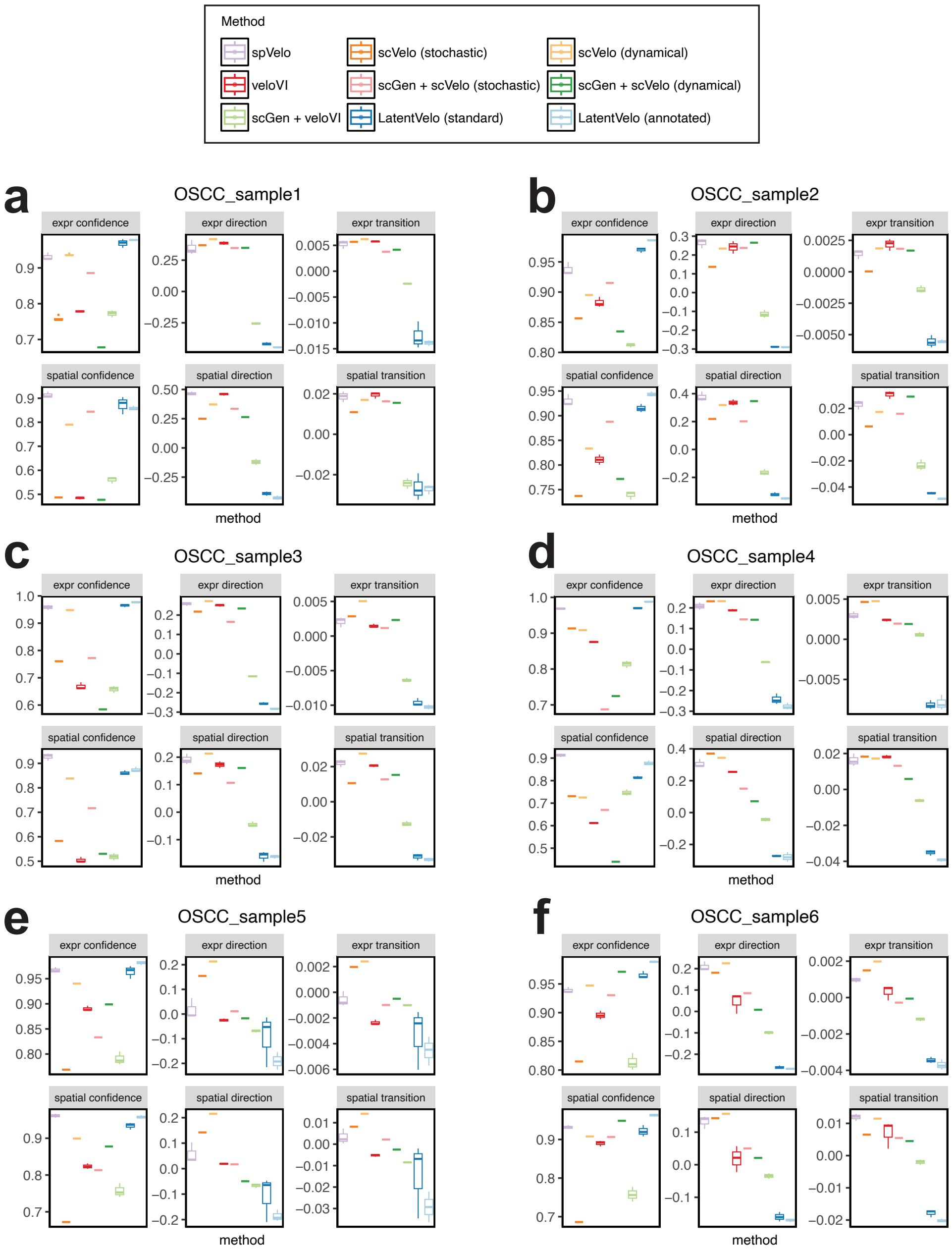


**Extended Data Fig. 5** Boxplots of all comparing scores in OSCC dataset. (a-f) Perbatch scores from sample 1 to sample 6 of OSCC dataset.

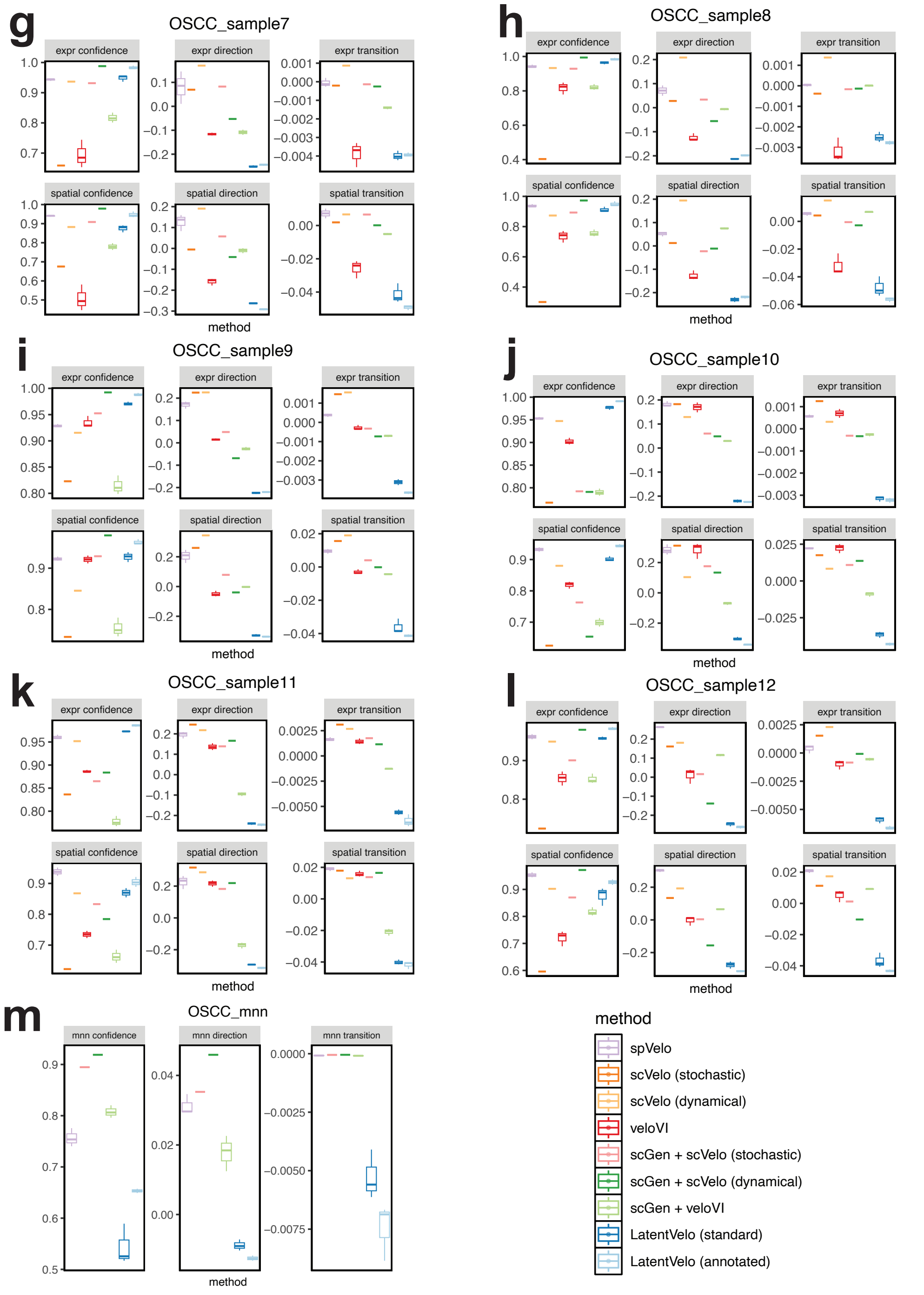


**Extended Data Fig. 5 (continue)** Boxplots of all comparing scores in OSCC dataset. (g-l) Per-batch scores from sample 7 to sample 12 of OSCC dataset. (m) MNN scores of OSCC dataset.

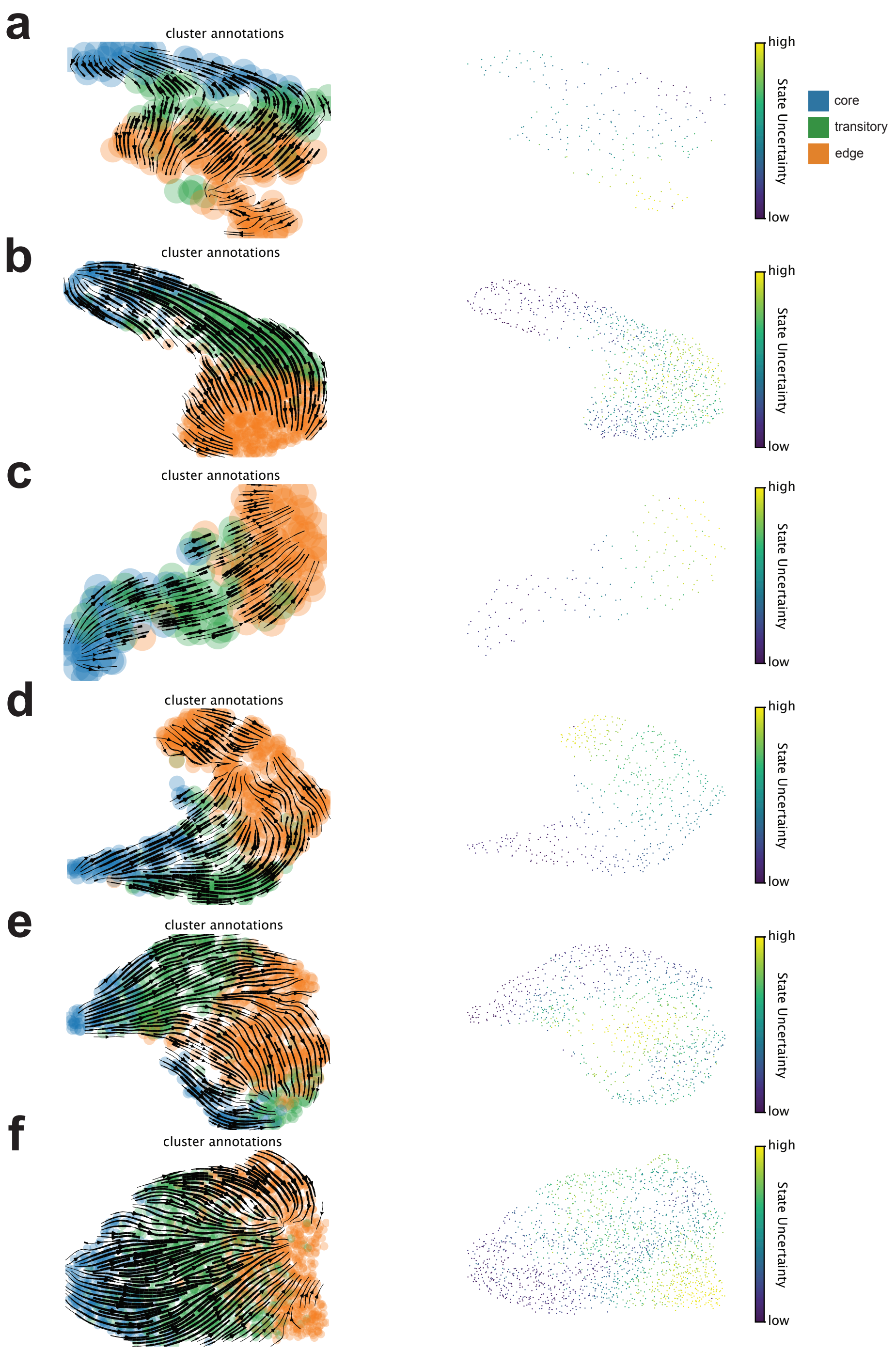


**Extended Data Fig. 6** Trajectory plots and uncertainty scatter plots of spVelo on OSCC dataset. (a-f) Trajectory plots on UMAP embedding from sample 1 to sample 6.

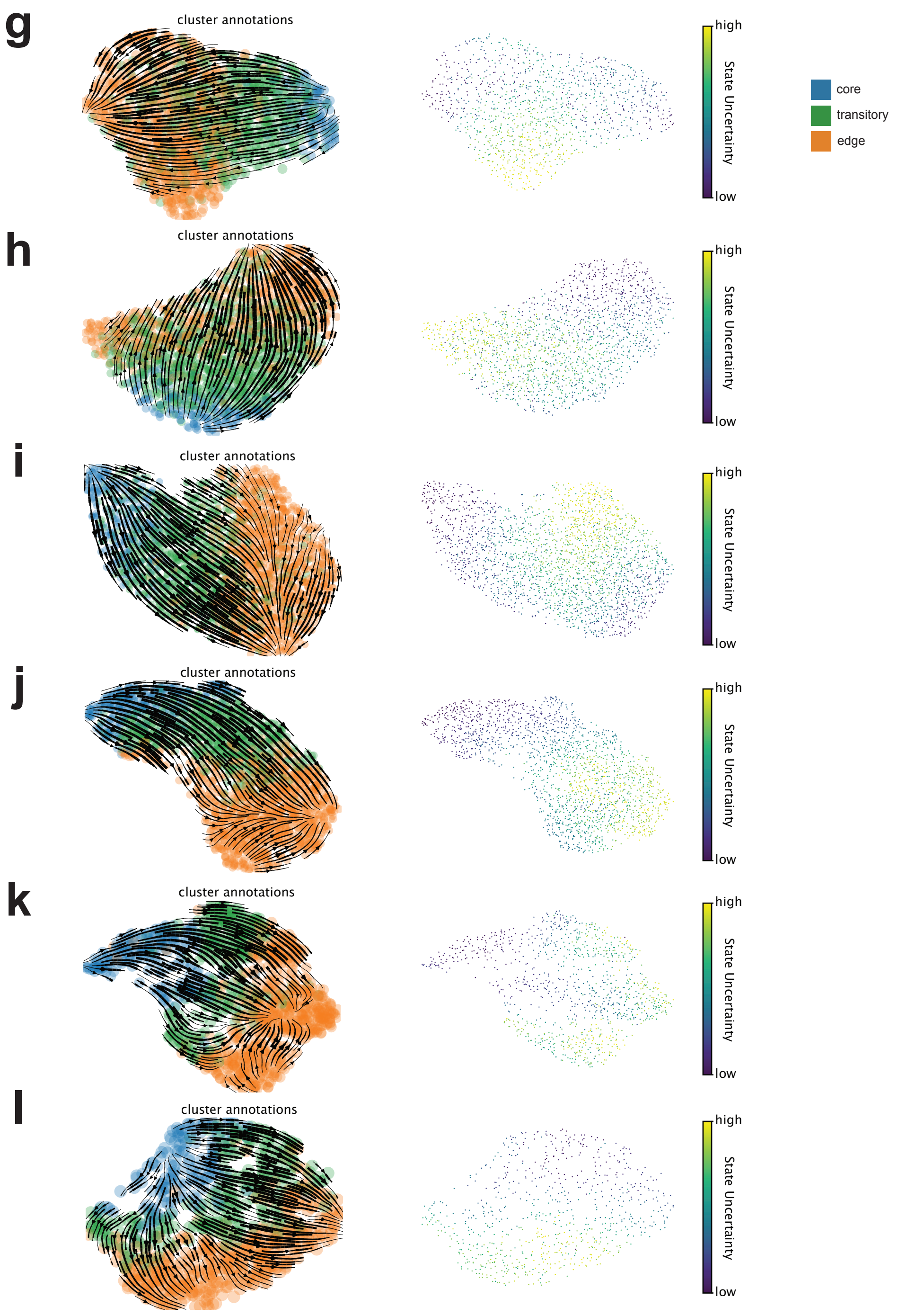


**Extended Data Fig. 6 (continue)** Trajectory plots and uncertainty scatter plots of spVelo on OSCC dataset. (g-l) Trajectory plots on UMAP embedding from sample 7 to sample 12.

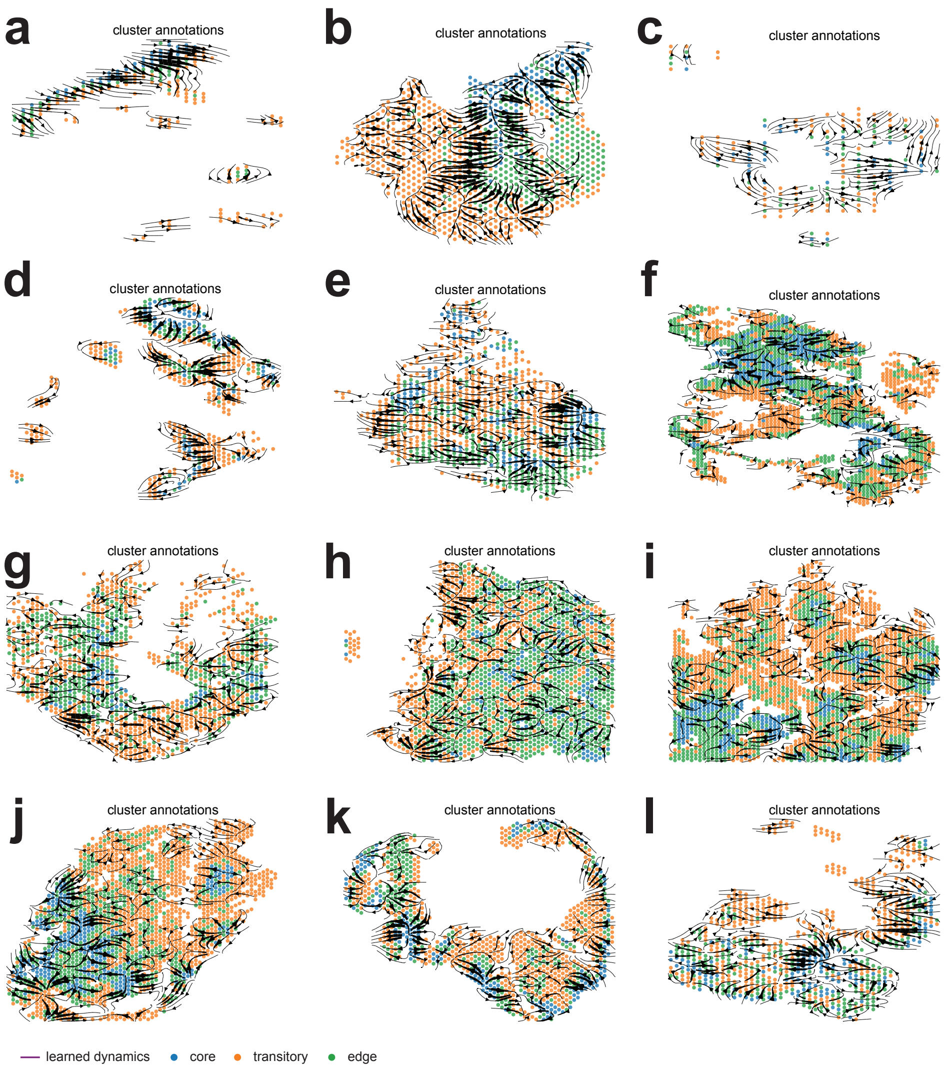


**Extended Data Fig. 7** Trajectory plots of spVelo on the spatial coordinate of OSCC dataset. (a-l) Trajectory plots on the spatial coordinate from sample 1 to sample 12.

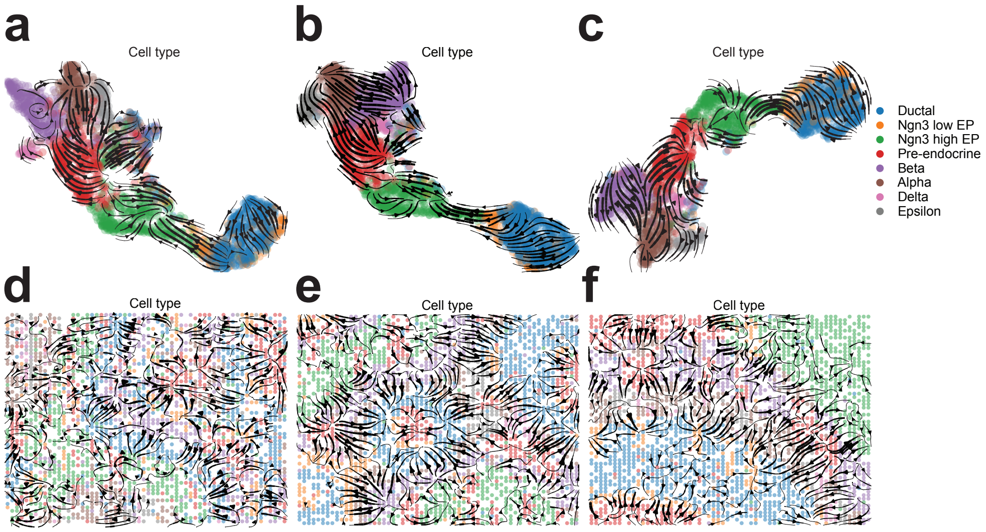


**Extended Data Fig. 8** Trajectory plots of spVelo on UMAP and spatial coordinate of simulated pancreas dataset. (a-c) Trajectory plots on UMAP embedding from batch 1 to batch 3. (d-f) Trajectory plots on spatial coordinate from batch 1 to batch 3.

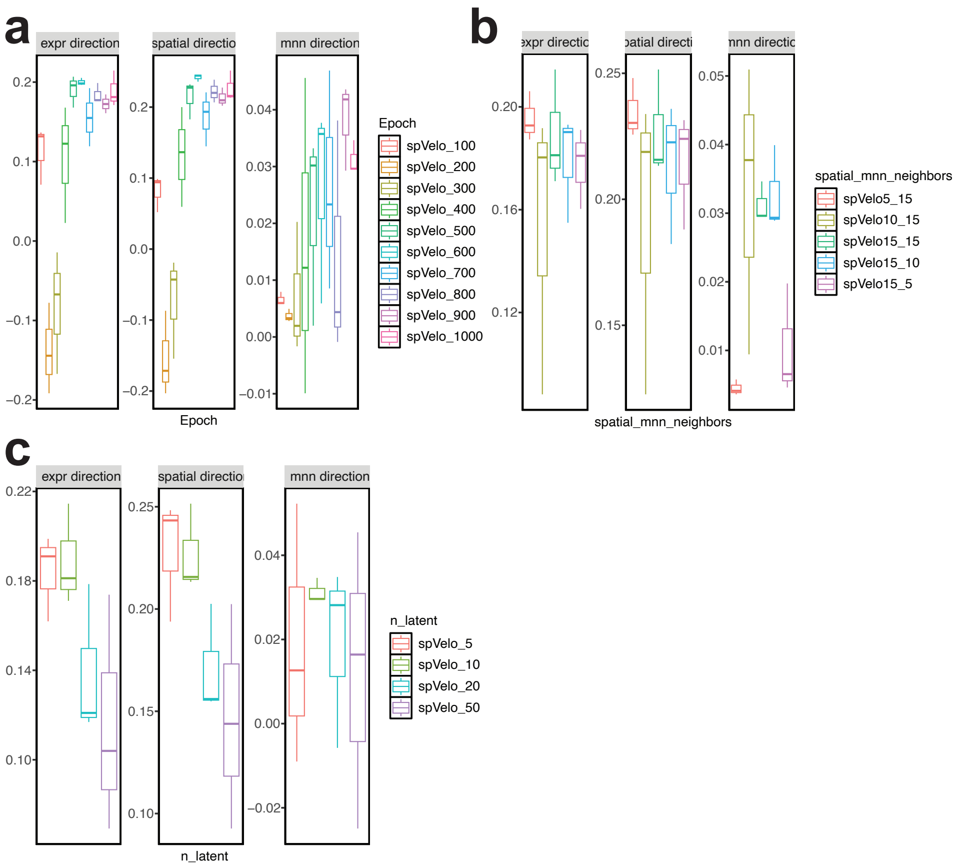


**Extended Data Fig. 9** Hyper-parameter tuning of spVelo on OSCC dataset. (a) The change of direction scores by adjusting the number of epochs. (b) The change of direction scores by adjusting the number of spatial and mnn neighbors. (c) The change of direction scores by adjusting the size of latent space.

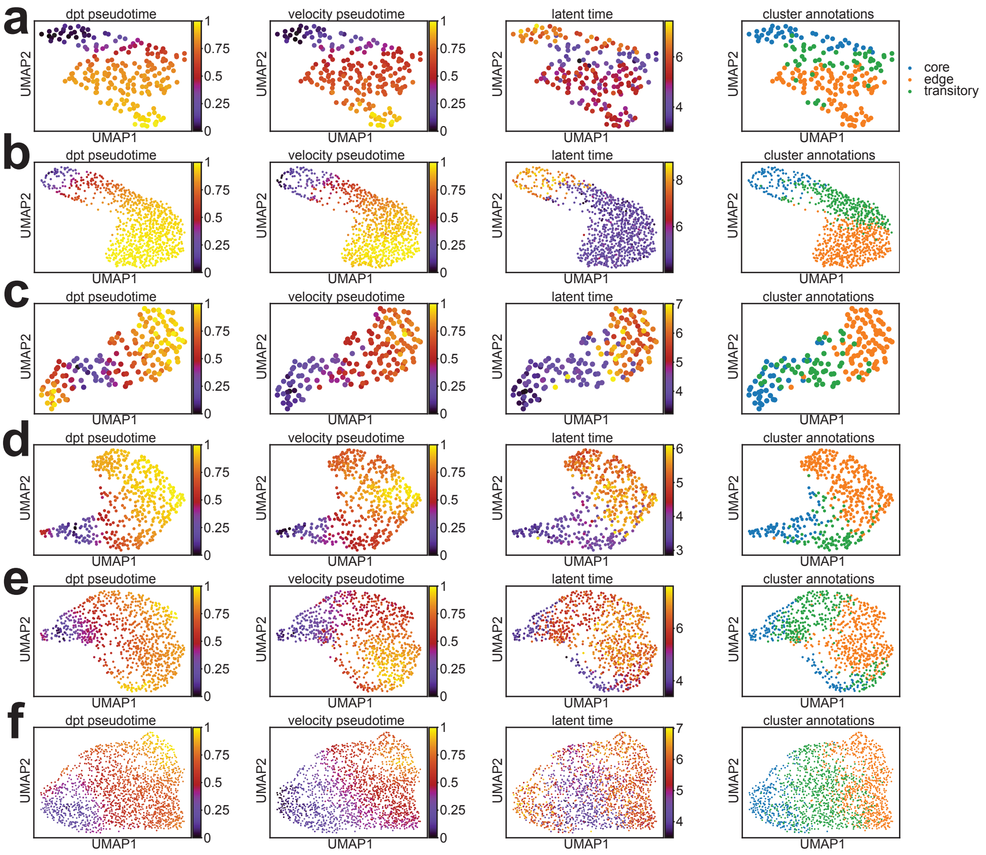


**Extended Data Fig. 10** Comparison of latent time inferred by spVelo with DPT and velocity pseudo-time. (a-f) Scatter plots from sample 1 to sample 6.

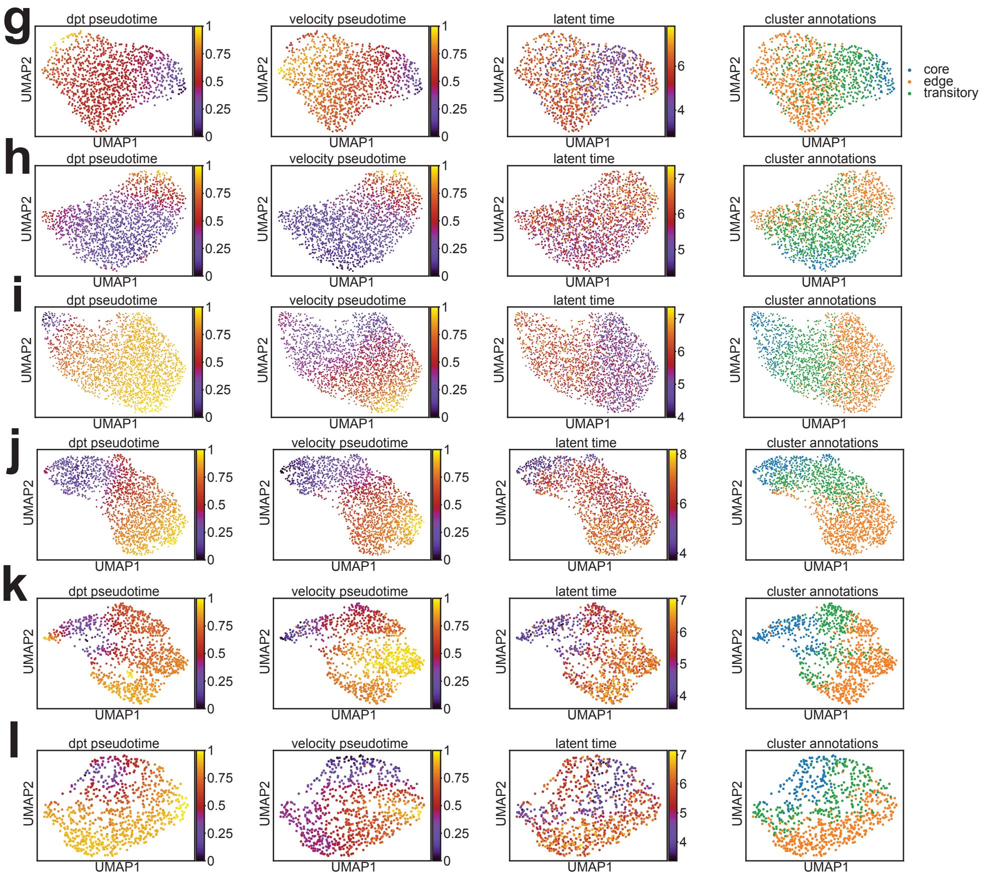


**Extended Data Fig. 10 (continue)** Comparison of latent time inferred by spVelo with DPT and velocity pseudo-time. (g-l) Scatter plots from sample 7 to sample 12.

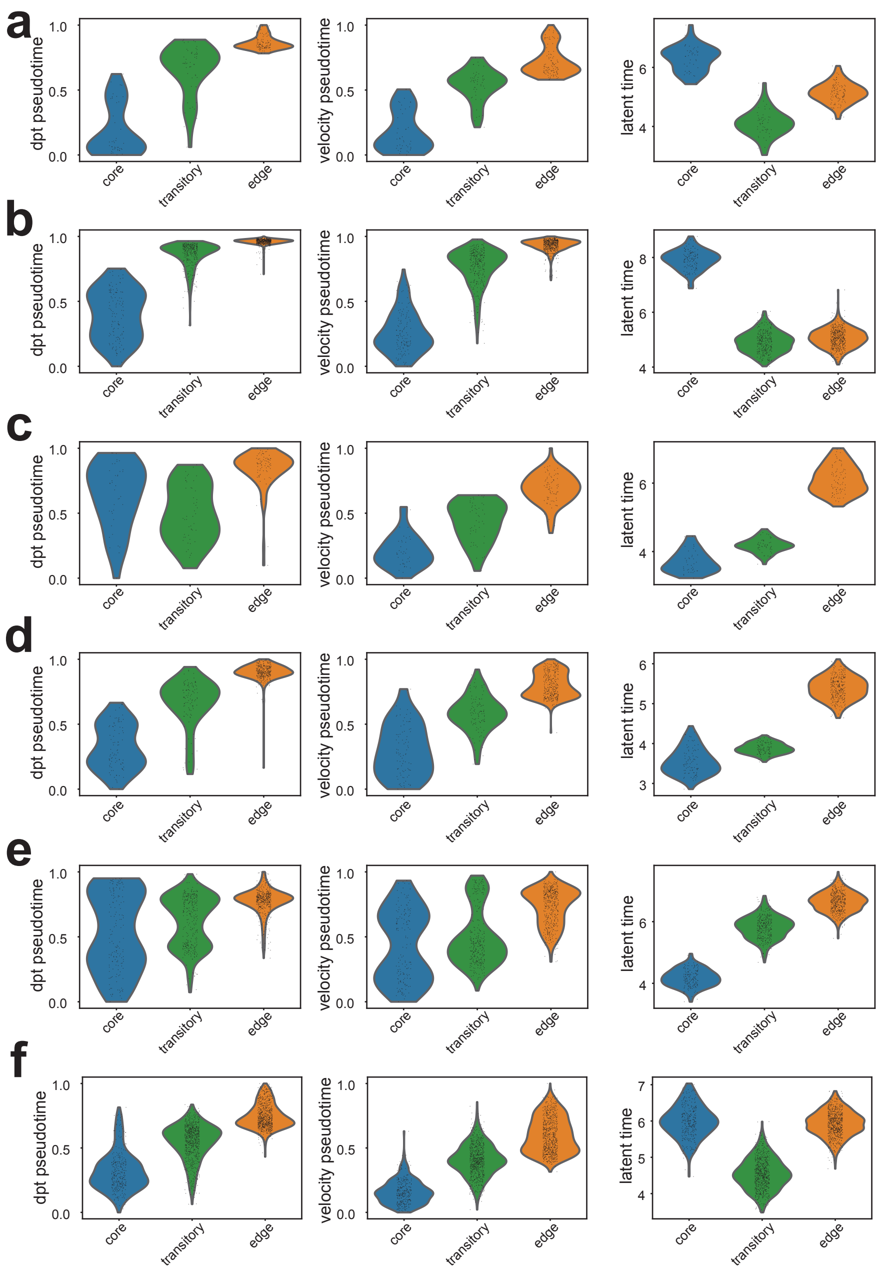


**Extended Data Fig. 10 (continue)** Comparison of latent time inferred by spVelo with DPT and velocity pseudo-time. (a-f) Violin plots from sample 1 to sample 6.

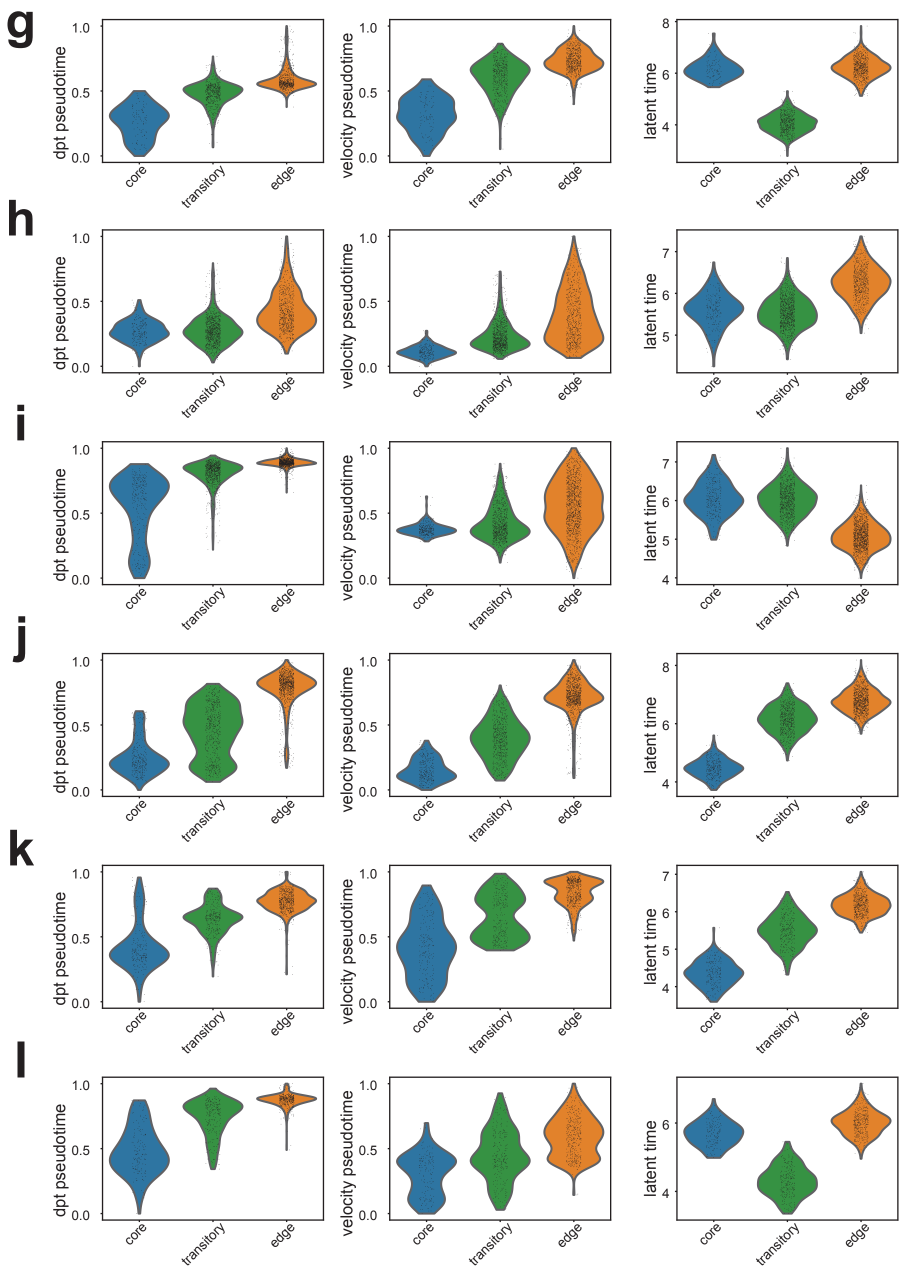


**Extended Data Fig. 10 (continue)** Comparison of latent time inferred by spVelo with DPT and velocity pseudo-time. (g-l) Violin plots from sample 7 to sample 12.

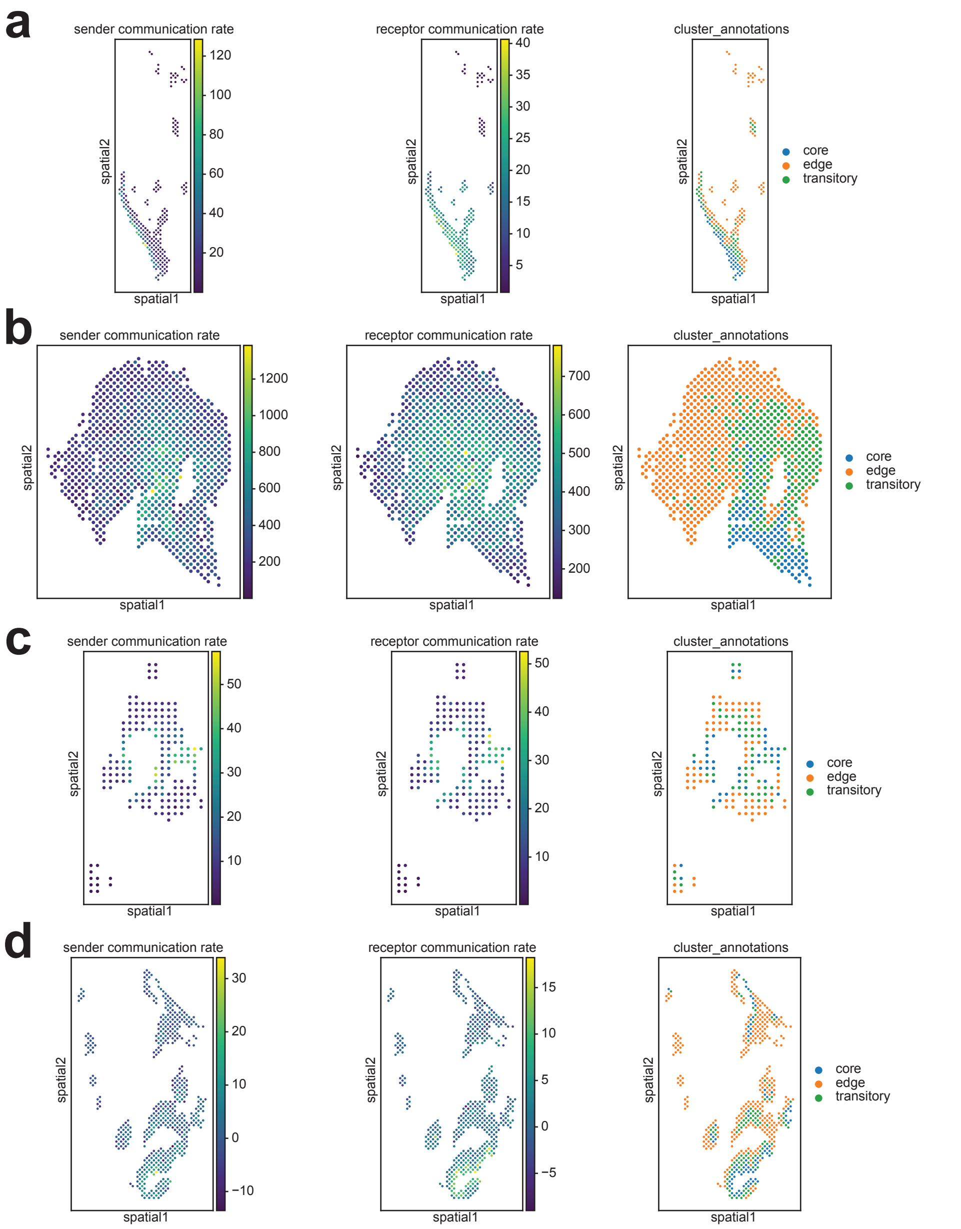


**Extended Data Fig. 11** Temporal cell-cell communications inference results of OSCC dataset. (a-d) Temporal cell-cell communications inference results of sample 1 to sample 4.

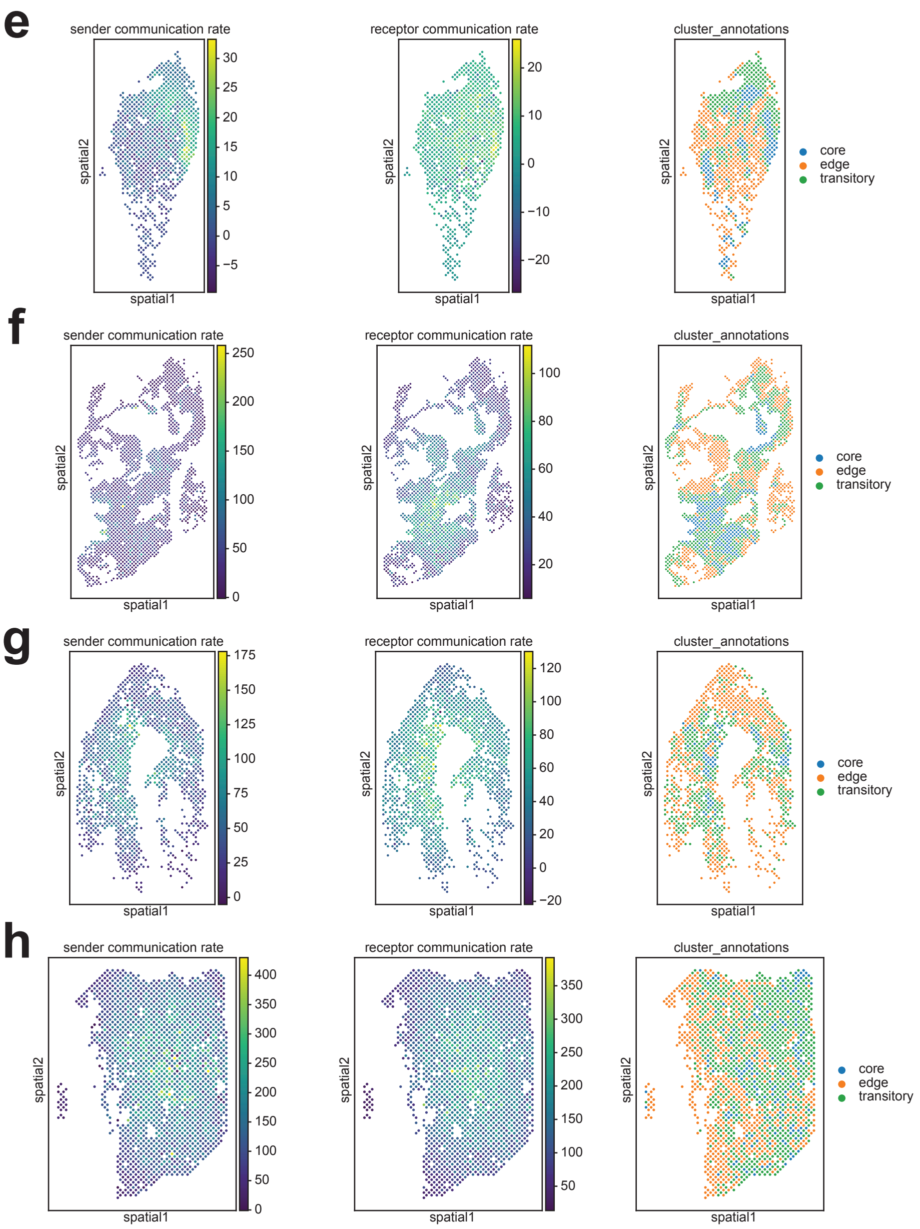


**Extended Data Fig. 11 (continue)** Temporal cell-cell communications inference results of OSCC dataset. (e-h) Temporal cell-cell communications inference results of sample 5 to sample 8.

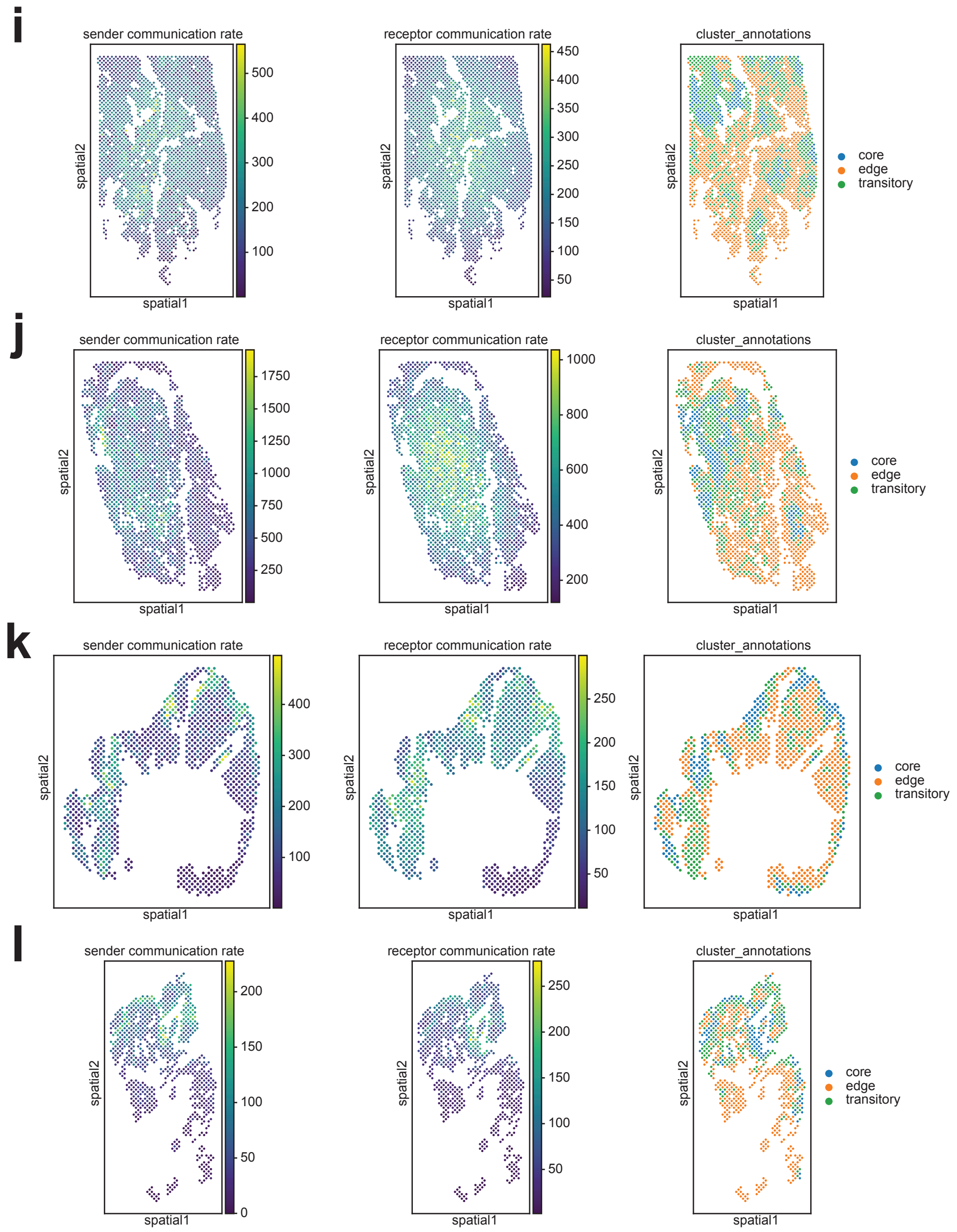


**Extended Data Fig. 11 (continue)** Temporal cell-cell communications inference results of OSCC dataset. (i-l) Temporal cell-cell communications inference results of sample 9 to sample 12.
